# High-Dose Paclitaxel and its Combination with CSF1R Inhibitor in Polymeric Micelles for Chemoimmunotherapy of Triple Negative Breast Cancer

**DOI:** 10.1101/2022.08.12.503695

**Authors:** Chaemin Lim, Duhyeong Hwang, Mostafa Yazdimamaghani, Hannah Marie Atkins, Hyesun Hyun, Yuseon Shin, Jacob D. Ramsey, Charles M. Perou, Marina Sokolsky-Papkov, Alexander V. Kabanov

**Author notes:** These authors contributed equally to this work (Chaemin Lim, Duhyeong Hwang). Corresponding Authors: Alexander V. Kabanov and Marina Sokolsky-Papkov.

## Abstract

The presence of immunosuppressive immune cells in cancer is a significant barrier to the generation of therapeutic immune responses. Similarly, *in vivo* triple-negative breast cancer (TNBC) models often contain prevalent tumor-associated macrophages in the tumor microenvironment (TME), resulting in breast cancer initiation, invasion, and metastasis by generating immunosuppressive environment. Here, we test systemic chemoimmunotherapy using small-molecule agents, paclitaxel (PTX), and colony-stimulating factor 1 receptor (CSF1R) inhibitor, PLX3397, to enhance the adaptive T cell immunity against TNBCs in immunocompetent mouse TNBC models. PTX and PLX3397 are very poorly soluble in water and shown poor therapeutic outcomes in TNBC animal models in conventional formulation. To address the challenge for the delivery of insoluble drugs to TNBC, we use high-capacity poly(2-oxazoline) (POx)-based polymeric micelles to greatly improve the solubility and widen the therapeutic index of such drugs. The results demonstrate that high-dose PTX in POx, even as a single agent, exerts strong effects on TME and induces the long-term immune memory. In addition, we demonstrate that the PTX and PLX3397 combination provides consistent therapeutic improvement across several TNBC models, resulting from the repolarization of the immunosuppressive TME and enhanced T cell immune response that suppress both the primary tumor growth and metastasis. Overall, the work emphasizes the benefit of drug reformulation and outlines potential translational path for both PTX and PTX with PLX3397 combination therapy using POx polymeric micelles for the treatment of TNBC.

## 1. Introduction

TNBC is characterized by low levels of expression of estrogen receptor, progesterone receptor, and human epidermal growth factor receptor 2 [1]. It is an aggressive disease that accounts for 15% to 20% of all breast cancers and is associated with poor prognosis and high recurrence rate, which annually contributes to ca. 5% of all cancer-related deaths [2]. TNBC is also known for aggressive metastasis to distant organs, that is encountered in over 40 % of TNBC patients [3]. In addition to relatively mixed response to traditional chemotherapies [1, 3], the immune checkpoint inhibitors have also relatively modest results in improving therapeutic outcomes and the management of the TNBC. In TNBC, the objective response rate for αPD-1 therapy is between 8-19%, with no durable clinical responses [4, 5]. However, when nab-paclitaxel (but not conventional paclitaxel) and Atezolizumab (Tecentriq) that targets PD-L1 were combined, this showed improved treatment outcomes, and initially obtained approval by the US Food and Drug Administration (FDA) as the new standard of care in patients with PD-L1 expression [6]. Unfortunately, Genentech, later voluntarily withdraw the U.S. accelerated approval for this treatment (https://www.gene.com/media/press-releases/14927/2021-08-27/genentech-provides-update-on-tecentriq-u). Therefore, developing, new effective strategies to improve TNBC immunotherapy is highly needed.

The tumor microenvironment (TME) is a critical component of tumor growth with immunosuppressive proteins and cell populations limiting the productive anti-tumor immune response [7–10]. In TNBC, the recruitment of immunosuppressive immune cells such as myeloid derived suppressor cells (MDSCs), tumor-associated macrophages (TAMs), and regulatory T cells (Tregs) and B cells (Bregs) by the tumor is a significant impediment to the generation of an effective immune response [11]. The most prevalent antigen-presenting cells within the TME are TAMs [12, 13]. They can promote breast cancer initiation, angiogenesis, invasion, and metastasis by generating an immunosuppressive TME via releasing cytokines, chemokines, and growth factors [14]. The presence of TAMs in TME is associated with poor clinical outcomes in TNBC [15–17]. Targeting the immunosuppressive TME has shown potential for improving immune checkpoint inhibitor therapy in patients [18].

The prevalence of TAMs in tumors and the potential for repolarizing to pro-inflammatory phenotype, which restores adaptive immune control of tumors, makes them an attractive target for establishing an immune-promoting TME [19]. Novel therapeutics aiming to target mechanisms promoting the survival, recruitment, polarization, and other properties associated with tumor-associated myeloid cells are currently in clinical development [20]. One strategy focuses on inhibition of the function and migration to the TME of immunosuppressive monocytes and macrophages. Colony-stimulating factor 1 (CSF-1) and its receptor, CSF-1R, play a major role in regulating proliferation and survival of macrophages and their precursors [21]. Inhibition of CSF1R with antibodies or small molecules suppresses breast cancer growth and reverses its resistance to chemo- and radiotherapy [22–24]. Blockade of macrophage recruitment with CSF1R-signaling antagonists, in combination with paclitaxel (PTX), enhanced CD8+ T cell immunity and improved survival of mammary tumor–bearing mice by slowing primary tumor development and reducing pulmonary metastasis via T cell–mechanism [22]. A number of small molecule inhibitors of CSF-1R (Pexidartinib, RXDX-105, BLZ945, Linifanib) are currently in the clinical trials for treatment of solid tumors.[20, 25] They show promise to attenuate immune escape of tumors and potentiate the effect of CPI immunotherapy and traditional cytotoxic therapy [13]. However, no intravenous (iv) formats for these agents are currently available to improve their delivery to tumor and metastatic sites, and to avoid dysphagia and other common complications of oral administration after chemo- and radiotherapy [26].

Here, we present systemic chemoimmunotherapy using small-molecule agents, PTX which is one of the most important therapeutic drugs in clinical management of TNBC [27] and CSF1R inhibitor, PLX3397 (Pexidartinib) that targets myeloid cells to enhance the adaptive T cell immunity. In a free form these agents are very poorly soluble in water and have poor distribution to the tumors. To address the challenge for the delivery of lipophilic drugs to tumors we use high-capacity POx-based polymeric micelles, a transformative technology, which is uniquely suited to greatly improve the solubility and widen therapeutic index of such drugs. This approach transforms a disadvantage of poorly soluble drugs into an advantage by packing them into small tumor-permeating polymeric micelle nanoparticles, improving their tumor distribution and enabling high dose therapy due to increased safety [28, 29]. Our findings suggest that high dose PTX and its co-loaded combination with PLX3397 enabled in POx micelles, remodel immunosuppressive TME, induce T cell-mediated antitumor response, produce immune memory and overall show promise in chemoimmunotherapy in animal models of primary and metastatic TNBC.

## 2. Results

### 2.1. Preparation and characterization of drug-loaded polymeric micelles

POx-based polymeric micelle formulations containing PTX and PLX3397 were prepared using a thin film method as previously reported [30]. In prior work we determined that POx block copolymers can efficiently solubilize PTX (POx-PTX), forming well-defined spherical micelles in aqueous media with extremely high drug loading and increased PTX solubility of up to 40 mg/mL [31]. In the current study, we solubilized PLX3397 in POx micelles (POx-PLX). As presented in **Figure 1(a)**, PLX3397 incorporated in the micelles with nearly no loss of the drug (98.8 % LE) and at extremely high drug loading (44.1 % LC). The overall solubility of PLX3397 in the micellar solution was also greatly increased to at least 20 mg/mL compared to the insolubility of the drug in water. The resulting POx-PLX micelles were spherical and extremely well-defined, with the particle size of 38 nm and PDI of 0.05, as determined by DLS. The PTX and PLX3397 combination micelles (POx-PTX/PLX) were also easily prepared by co-loading these drugs at a 1/1 (w/w) drug ratio in the POx micelles (**Figure 1(a)**). There was little or no loss of the drugs upon loading (94.0 % LE for PTX and 99.9 % LE for PLX3397, respectively) and the resulting micelles also contained high amounts of drugs per polymer (the combined LC was as much as 44%; 21.2 % for PTX and 22.5 % for PLX3397). Like the single drug micelles, the co-loaded micelles were also spherical and well-defined (57 nm, PDI = 0.16).

**Figure 1.**
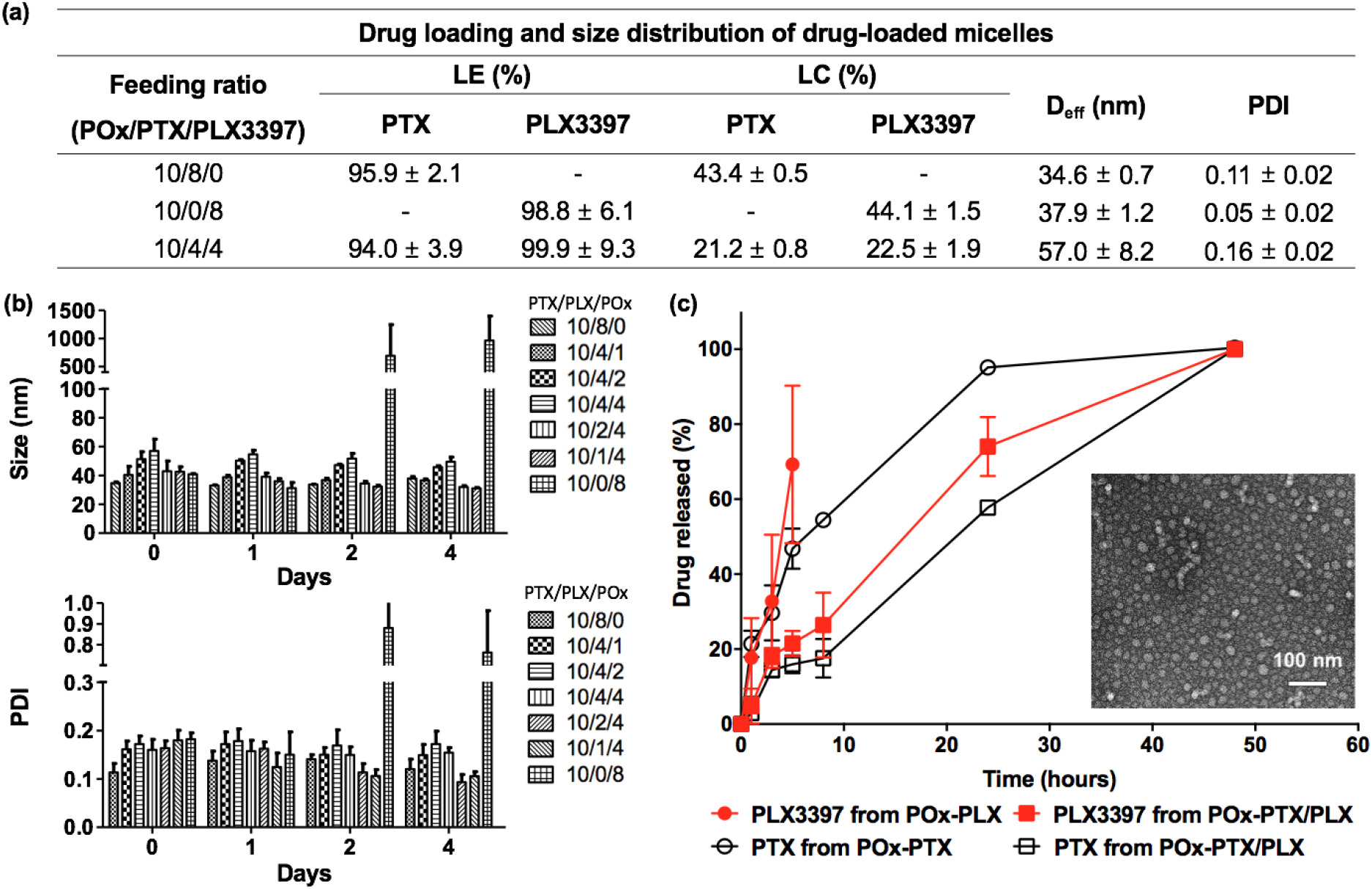
Physicochemical properties of drug-loaded micelles. (a) drug loading (expressed as loading efficiency (LE) (%) and loading capacity (LC) (%)) and size distribution (expressed as Deff and polydispersity index (PDI)) of drug-loaded micelles. (LE (%) = M_drug_ / M_drug added_ × 100 (%) and LC (%) = M_drug_ / (M_drug_ + M_excipient_) × 100 (%)) (b) stability of drug-loaded micelles in solution over time measured by dynamic light scattering (DLS). (c) drug release profiles of PLX3397 and PTX from either single (POx-PLX and POx-PTX, respectively) or POx-PTX/PLX and morphology of POx-PTX/PLX by transmission electron microscopy (TEM) (scale bar = 100 nm). All measurements were done triplicate and values are average of three replications with standard deviation except for TEM image.

For the stability studies the micelle solutions were kept at 4 °C for up to 4 days and then the particle size and PDI values were measured at room temperature by DLS. The single drug POx-PLX micelles aggregated at day 2, which was accompanied by drastic increase in the particle size and PDI, as well as visual drug precipitation. In contrast, the combination micelles retained their small size and PDI for the entire period of observation, with no drug precipitation observed (**Figure 1(b)**). The stabilization effect was obviously due to the contribution of the PTX as the single drug POx-PTX micelles are stable in solution for at least two weeks [32]. In any case the best strategy for these micellar formulations of the drugs is to keep them in the dry lyophilized form with rehydration prior to the use (**Figure S1**).

The drug release profiles were examined under sink conditions. The single drug POx-PTX micelles showed sustained drug release, with 50 % drug released at 5 h (**Figure 1(c)**). The single drug POx-PLX micelles also showed similar sustained drug release profile, however, precipitation was observed after 5 h. POx-PTX/PLX remained stable during entire observation time. Interestingly the release of each drug from the co-formulated POx-PTX/PLX micelles was slower than that from the single drug micelles, which could be attributed to attractive drug-drug/drug-polymer interactions in these complex mixtures (**Figure 1(c)**).

### 2.2 Drug combination micelles exhibit synergy in breast cancer cells

While the main objective of this study using PLX3397 was to deplete the macrophage population in TME to treat TNBC tumors, we point out that PLX3397 is also known as an inhibitor of c-Kit, a receptor tyrosine kinase, with half-maximal inhibitory concentration (IC_50_) of 10 nM [33, 34]. Therefore, this drug can elicit direct anti-cancer effect by inhibiting proliferation of tumor cells via c-Kit pathway, as was observed in many tumor models [35, 36]. Thus, we evaluated the cytotoxicity of the single and combination drug micelles in 4T1, T11-apobec and T12 TNBC mouse models cell line transplant tumors (**Figure 2 (a)** and **(b)**). In 4T1 cell line, the IC_50_ values of POx-PTX and POx-PLX were 25.9 *μ*g/ml and 151.3 *μ*g/ml, respectively. However, the POx-PTX/PLX displayed much lower IC_50_ values (for instance 0.72 *μ*g/ml of total drug at 1:1 mass ratio), suggesting strong synergistic effect between the drugs. A similar phenomenon was seen in the two other TNBC cell lines. For example, in T11-apobec the IC_50_ value for drug combination at 1:1 mass ratio was 0.02 *μ*g/ml compared to 0.09 *μ*g/ml and 1.06 *μ*g/ml for POx-PTX and POx-PLX, respectively. Therefore, we determined the combination indices (CI) of the PTX and PLX3397 in these cell models as defined by Chou and Talalay [37]. As very strong synergy between the drugs (CI ≪ 1) was observed in 4T1 and T11-apobec cells, for nearly entire range of the cell fraction affected (Fa) (**Figure 2(b)**). In the T12 cells there was some antagonism at Fa < 0.3 but overall, the drugs also displayed strong synergy at Fa > 0.3. In this cell line the IC_50_ value for the drug combination was still much lower that these for the single drug micelles (0.09 μg/ml for the combination at 1:1 drug ratio vs. 0.33 μg/ml for POx-PTX, and 0.76 μg/ml for POx-PLX). Interestingly, the drug synergy was exhibited at various PTX: PLX3397 drug mass ratios ranging from 4:1 to 1:4 as demonstrated using 4T1 cell model. The combination drug micelles also induced greater extent of apoptosis and necrosis in the TNBC cells (at equivalent drug concentration 1 μg/ml) as determined after 24-h exposure (**Figure 2(c)**). Overall, in the studied breast cancer models, the drug combinations were substantially more active than either of the single drugs. Based on this *in vitro* analysis, we fixed the POx to drug mass ratios at 10/4 for single drug micelles or 10/4/4 for POx-PTX/PLX micelles and proceeded to *in vivo* evaluation.

**Figure 2.**
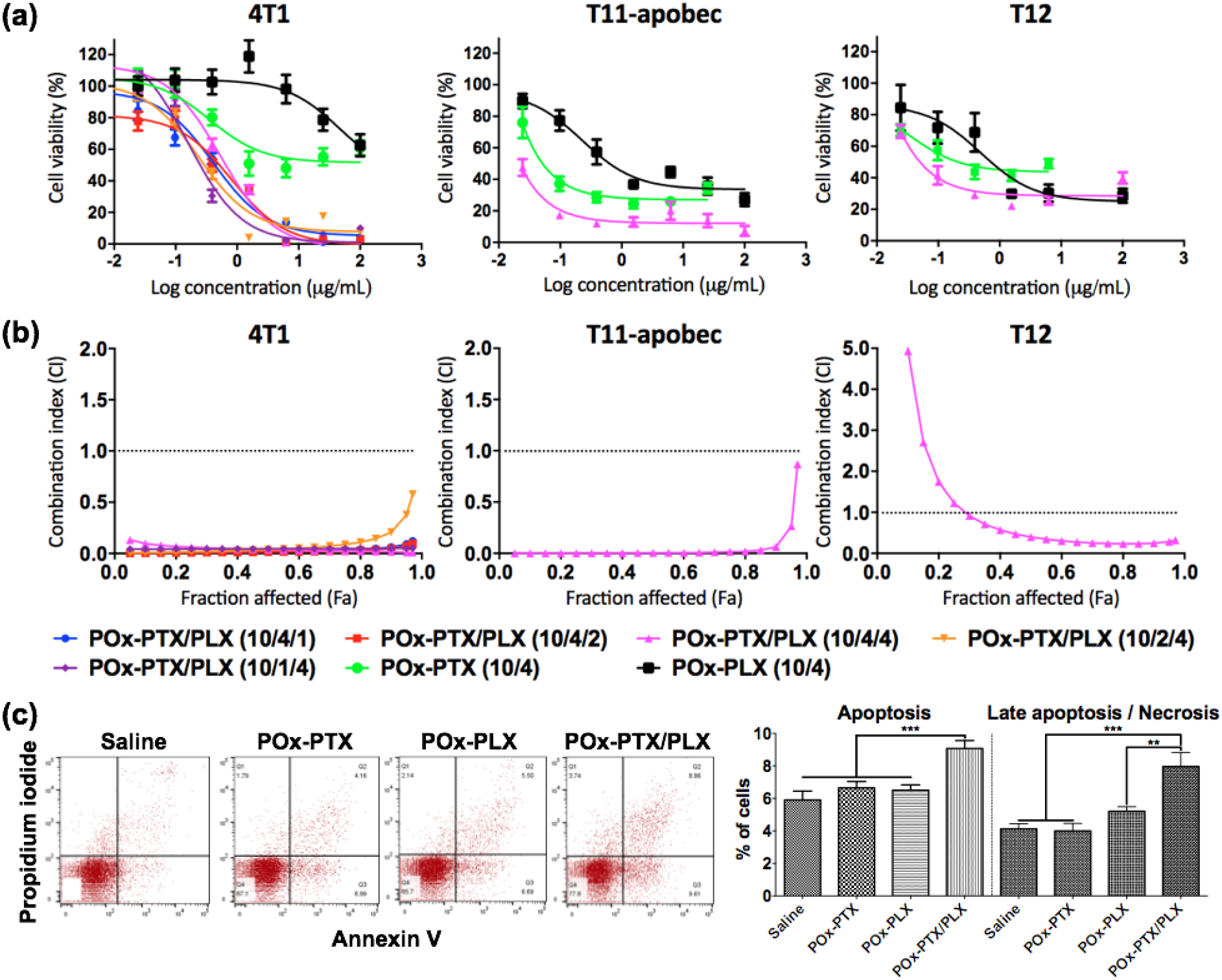
*in vitro* evaluation. (a) Cytotoxicity and (b) Combination index of drug-loaded micelles in the 4T1, T11-apobec and T12 cell lines. (a) Ordinate presents the logarithmic scales of the concentrations of the respective single drugs in POx-PTX or POx-PLX, or PTX in POx-PTX/PLX. (a, b) The numbers in the brackets designate the mass ratios POx: PTX (10/4) or POx: PLX3397 (10/4) for the single drug micelles, or POx: PTX: PLX3397 (10/4/1; 10/4/2; 10/2/4; 10/1/4) for the co-loaded micelles. (c) Flow cytometry analysis of apoptosis and necrosis using annexin V and propidium iodide double staining in 4T1 cells treated by saline, POx-PTX (10/4), POx-PLX (10/4) or (POx-PTX/PLX (10/4/4) at PTX and/or PLX3397 = 1 μg/ml. Statistical comparison was done using a one-way analysis of variance (ANOVA) and followed by Bonferroni post-tests for multiple comparison (n = 4~5). Statistical difference: *p < 0.05, **p < 0.01, *** p < 0.001.

### 2.3 Anti-tumor effects in TBNC models

We further explored antitumor activity of our agents in three immune-competent orthotopic TNBC models: (1) 4T1, (2) T11-apobec, and (3) T12 (**Figure 3(a-c)**). The 4T1 is an aggressive tumor that produces spontaneously metastases to the lungs, liver, lymph nodes, and brain [38] and is resistant to αPD-L1 therapy [39] but sensitive to radiation and chemotherapy [40, 41]. The two other tumors are derived from a TP53-/- genetically engineered mouse models (GEMMs) that have shown genomic and genetic similarities to human TNBCs [42–44]. T11-apobec has deletion in P53, overexpresses APOBEC3 to increase TMB, and closely recapitulates the genetic lesions and immune responses found in patients with claudin-low breast cancer [45]. The T12 tumor is another closely related claudin-low model that has high level of expression of CSF1R, and markers related to immunosuppressive TAMs [46]. It has previously shown a partial response to oral PLX3397 [46]. Prior to mouse treatments experiments we analyzed drug toxicity in various formulation to ensure that the doses used are safe (supplementary **Figure S2**).

**Figure 3.**
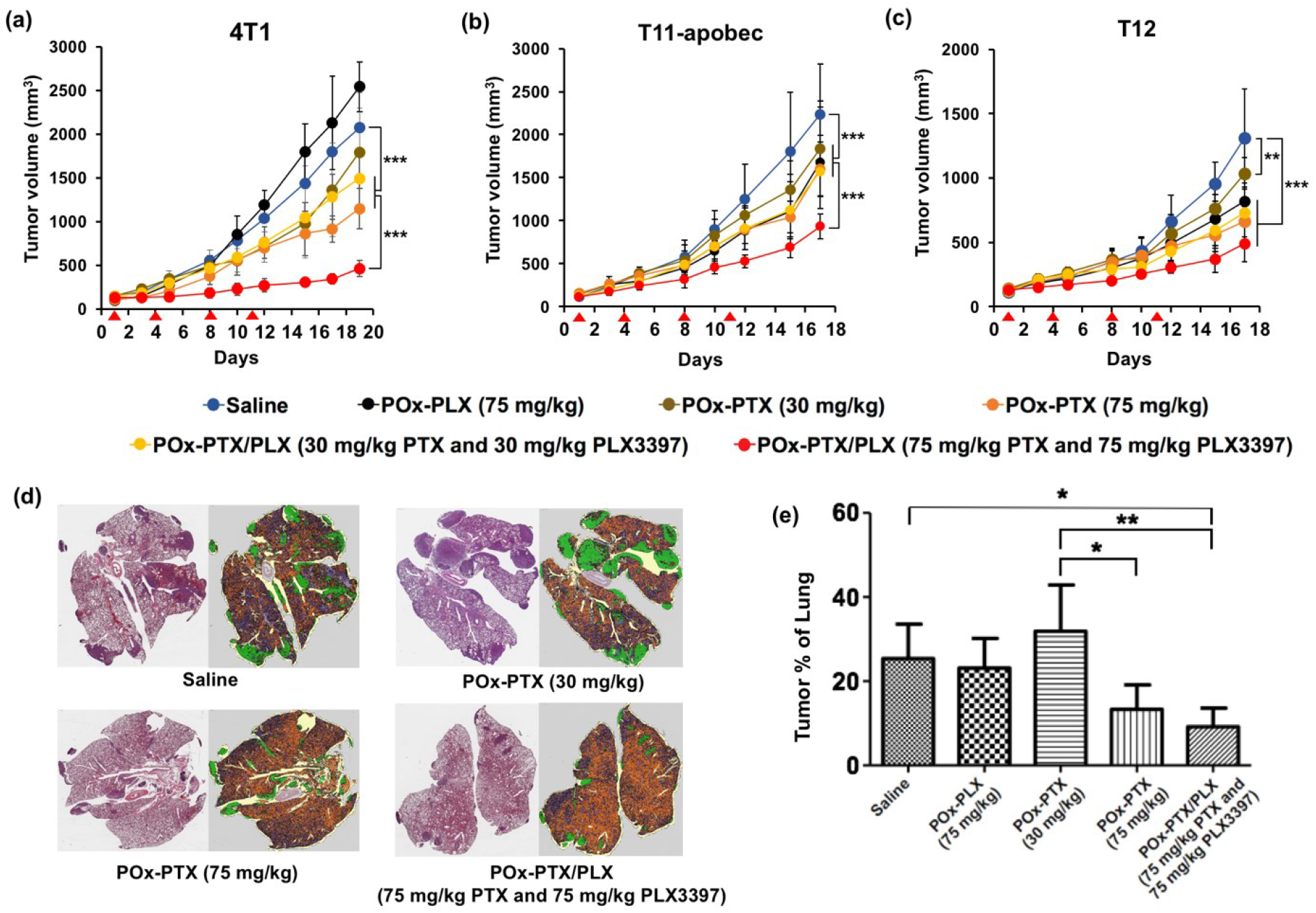
(a-c) Primary tumor inhibition and (d, e) suppression of lung metastases in mice bearing orthotopic TNBC tumors (a, d, e) 4T1 (b) T11-apobec (c) T12 (a-c). The animals (n = 4~8) were injected iv (as shown by arrows, q4d × 4) with: saline, POx-PLX (75 mg/kg), POx-PTX (30 mg/kg), POx-PTX (75 mg/kg), POx-PTX/PLX (30 mg/kg PTX and 30 mg/kg PLX3397) and POx-PTX/PLX (75 mg/kg PTX and 75 mg/kg PLX3397). **(d)** Representative H&E and overlay images of lungs from mice with 4T11 tumors harvested between 24 days, and **(e)** quantification of lung metastasis following indicated treatments. Statistical comparison of data for tumor inhibition (a-c), and metastases (e) was done using a two-way ANOVA and followed by Bonferroni post-tests for multiple comparison (n = 4~5). Statistical difference: * (p < 0.05), ** (p < 0.01), and *** (p < 0.001). See supplementary **Table S1** for the complete statistical comparison between all groups for primary tumors (a-c).

#### 4T1 orthotopic model of TNBC

The tumor growth curves are presented in **Figure 3(a)**. In the saline control group, the tumor volume increased from ca. 100 to ca. 2000 mm^3^ over 20 days. In this TNBC model neither POx-PLX (75 mg/kg) nor POx-PTX (30 mg/kg) were able to slow down the tumor growth. The groups treated with the high-dose POx-PTX (75 mg/kg) displayed significant tumor inhibition effects compared to the saline control or POx-PTX (30 mg/kg). The co-loaded POx-PTX/PLX produced the most pronounced antitumor effect that was also dose-dependent (**Figure 3(a)**, and supplementary **Figure S3**). The treatment with the higher dose POx-PTX/PLX (75 mg/kg PTX and 75 mg/kg PLX3397) surpassed all other treatments including POx-PTX (75 mg/kg). The POx-PTX/PLX at this dose also showed superiority vs. the combination treatment with the maximum tolerated dose (MTD) of PTX and PLX3397 in the standard Cremophor EL vehicle (Supplementary **Figure S3**) and increased apoptosis in the tumor tissue (Supplementary **Figure S4**). Along with the effect on the primary tumor, both the high-dose POx-PTX and POx-PTX/PLX significantly decreased the levels of metastases in lung tissues compared to the POx-PLX, lower dose POx-PTX (30 mg/kg), and the saline control (**Figure 3(d) and 3(e)**).

#### T11-apobec orthotopic model of TNBC

The combination treatment with POx-PTX/PLX (75 mg/kg PTX and 75 mg/kg PLX3397) also showed significant anti-tumor effect in the orthotopic T11-apobec model compared to the saline or other treatment groups (**Figure 3(b)**). The antitumor effects of single drugs or drug combination at the lower dose were significant compared to the saline control but small and indistinguishable from each other. (**Figure 3(b)**).

#### T12 orthotopic model of TNBC

In this model each single drug-loaded POx micelle treatments caused modest tumor inhibition. Like in the previous models the POx-PTX/PLX (75 mg/kg PTX and 75 mg/kg PLX3397) exhibited most pronounced antitumor effect, but differences were not statistically significant compared to POx-PTX (75 mg/kg) (**Figure 3(c)**).

### 2.4 Effect of drug co-formulation vs. separate administration

We further explored whether there is a benefit in co-loading PTX and PLX3397 into a single micellar vehicle vs. administering them separately (**Figure 4(a)**). We compared two drugs in one micelle, POx-PTX/PLX, with the same drugs loaded in separate micelles and mixed immediately prior to injection or administered sequentially. Since PLX3397 is orally available [36], we also explored the combination treatment with POx-PTX micelles injected iv and free PLX3397 given by oral gavage in a standard vehicle (5% DMSO, 45% PEG300, and 5% Tween 80 in distilled water) using previously described dose regimen [47]. As a reference point, we also included a group treated with POx-PTX. The results of this experiment suggest a clear benefit of co-loaded drug micelles compared to separate treatments (**Figure 4(b)**). Although the combination treatment with POX-PTX iv and PLX3397 orally showed some improvement over single POx-PTX, this combination was much less effective compared to POx-PTX/PLX at the same cumulative dose. Likewise, the sequential iv administration of POx-PTX and POx-PLX was more active that POx-PTX, but less active than co-loaded POx-PTX/PLX. The separate micellar drugs mixed prior to injection have shown similar activity to that of POx-PTX/PLX, which could be explained by the rapid inter-micellar drug exchange resulting in reconstitution of the co-loaded system [28, 29, 48].

**Figure 4.**
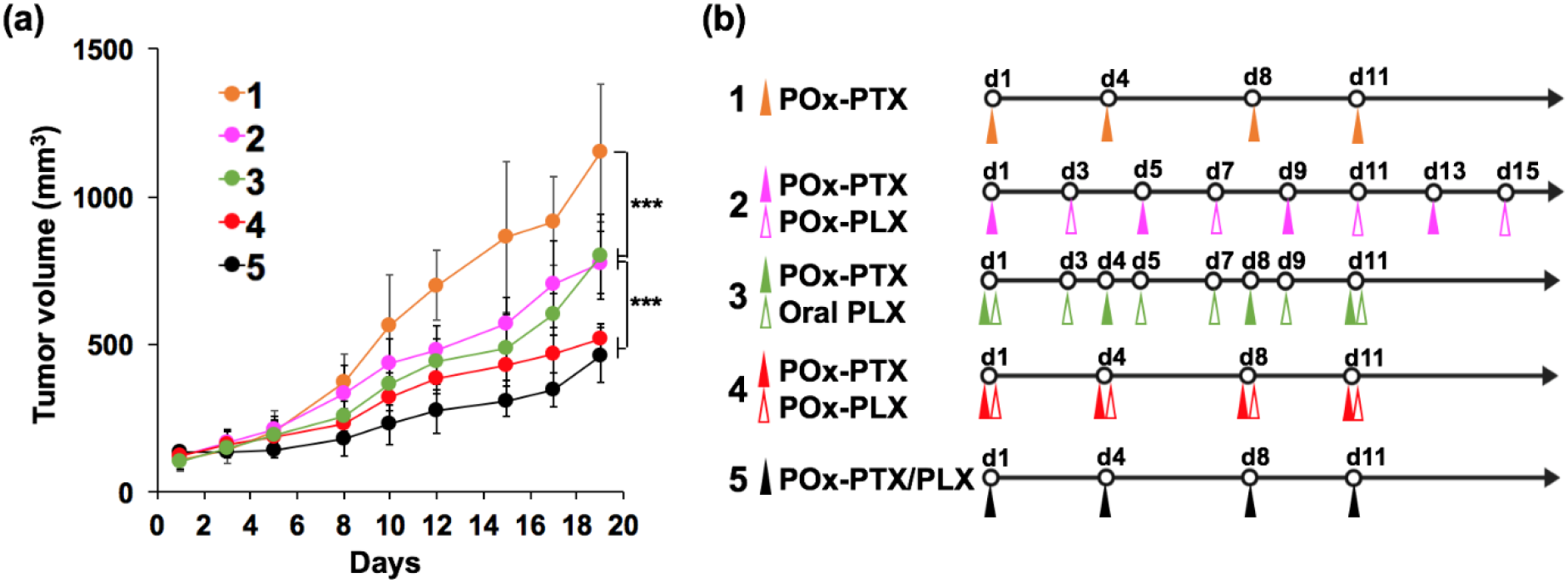
Dependence of anti-tumor effects of the PTX and PLX3397 combination therapy on the drug formulation, dose regimen and administration route in mice with orthotopic TNBC 4T1 tumors. **(a)** Antitumor effects in animals with tumors treated as schematically shown in **(b)** with 1 – POx-PTX (75 mg/kg), 2 – sequentially injected POx-PTX (75 mg/kg) and POx-PLX (75 mg/kg), 3 – POx-PTX (75 mg/kg) and oral PLX3397 (50 mg/kg) in standard vehicle, 4 – POx-PTX (75 mg/kg) and POx-PLX (75 mg/kg) mixed immediately prior to injection (simultaneous), 5 – co-loaded POx-PTX/PLX (75 mg/kg PTX and 75 mg/kg PLX3397). All treatment groups received drugs iv except for 3 where the micellar POx-PTX was injected iv and PLX3397 was administered orally. The 1 and 2 datasets are same as in **Figure 3(a)**. Statistical comparison of data for tumor inhibition was done using a two-way ANOVA and followed by Bonferroni post-tests for multiple comparison (n = 4~5). Statistical difference: *** p < 0.001. See supplementary **Table S3** for the compete statistical comparison between all groups.

### 2.5 Assessment of activation of immune cells population in TME

To uncover effects of our treatments on the TME we harvested *in vivo* tumors and performed immune phenotyping by multi-panel parameter flow cytometry (**Figure 5**, and supplementary **Figure S5**, **Table S4,** and **S5**). In the 4T1 model, flow cytometry showed that CD8^+^ T cells and CD4^+^ T cells were markedly increased in all PTX containing treatment groups compared to control or PLX3397 treated groups, while there were no statistical differences between the low and high dose POx-PTX and/or POx-PTX/PLX treated groups. In the T12 orthotopic model, similar trends were observed for CD8^+^ T cells and CD4^+^ T cells, but differences were not statistically significant in the group comparisons (1-way ANOVA with Turkey post-test, see Supplementary **Table S6**). In the T11-apobec model, the POx-drug formulation treatment did not change the CD8^+^ populations compared to the control group, but CD4^+^ were decreased by the POx-PTX/PLX treatment. There were no statistical changes in Tregs population after the treatments compared to control groups in any of the three tumor models.

**Figure 5.**
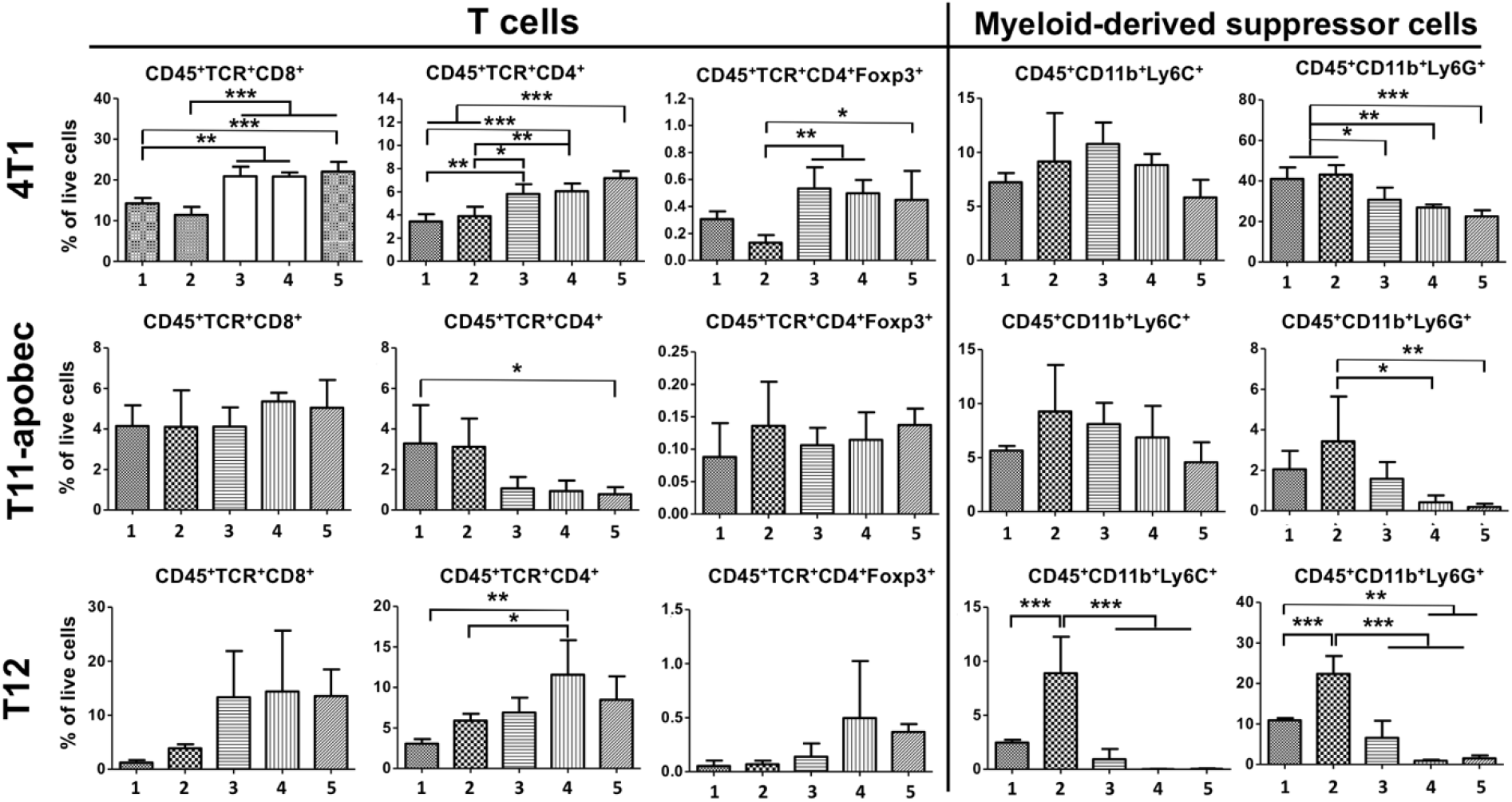
Immune phenotyping to demonstrate the effect of drug formulations on TNBC tumor models. The animals received saline or drug-loaded micelles iv using q4d × 2 with 1 – normal saline, 2 –POx-PLX (75 mg/kg), 3 – POx-PTX (30 mg/kg), 4 – POx-PTX (75 mg/kg), and 5 – co-loaded POx-PTX/PLX (75 mg/kg PTX and 75 mg/kg PLX3397). 3 Days after 2^nd^ dose of treatment, tumors were harvested to perform flow cytometry to show the impact on CD8^+^ T cells, CD4^+^ T cells, Treg, M-MDSCs and G-MDSCs. Statistical comparison of data for tumor inhibition was done using a one-way ANOVA with Tukey’s test for multiple comparison (* (p < 0.05), ** (p < 0.01), and *** (p < 0.001)). (see Supplementary **Table S6** for statistical comparisons between all groups)

We also assessed the MDSC subpopulations (**Figure 5**). In 4T1 or T11-apobec models POx-PLX had no effect on granulocytic MDSC (G-MDSC) or monocytic MDSC (M-MDSC). Interestingly, in T12 model, POx-PLX increased both M-MDSC and G-MDSC subpopulations. while POx-PTX and POx-PTX/PLX depleted these subpopulations. Both POx-PTX and POx-PTX/PLX depleted G-MDSC and produced no changes on M-MDSC in 4T1 or T11-apobec tumors. Statistical analysis showed a trend or significant difference in G-MDSC between POx-PTX (30 mg/kg) > POx-PTX (75 mg/kg) > POx-PTX/PLX groups (see Supplementary **Table S6**).

Finally, we assessed TAMs (**Figure 6**). As expected, POx-PLX decreased the total TAMs in all three tumor models. Interestingly, this drug enhanced the M1-like macrophages in 4T1 and T11-apobec but not in T12 tumors, while M2-like macrophages were not changed in any of these tumors. The difference in response may be due to the enrichment of T12 tumors with M1-like macrophages [46]. However, in all models POx-PLX decreased the M2/M1 ratios suggesting pro-inflammatory repolarization of the TAMs. Interestingly, POx-PTX also appeared to decrease the M2/M1 ratio at least in the case of 4T1 tumors (see Supplementary **Table S6** for analysis of significance). The POx-PTX/PLX combination decreased the M2/M1 ratios compared to controls in all cases.

**Figure 6.**
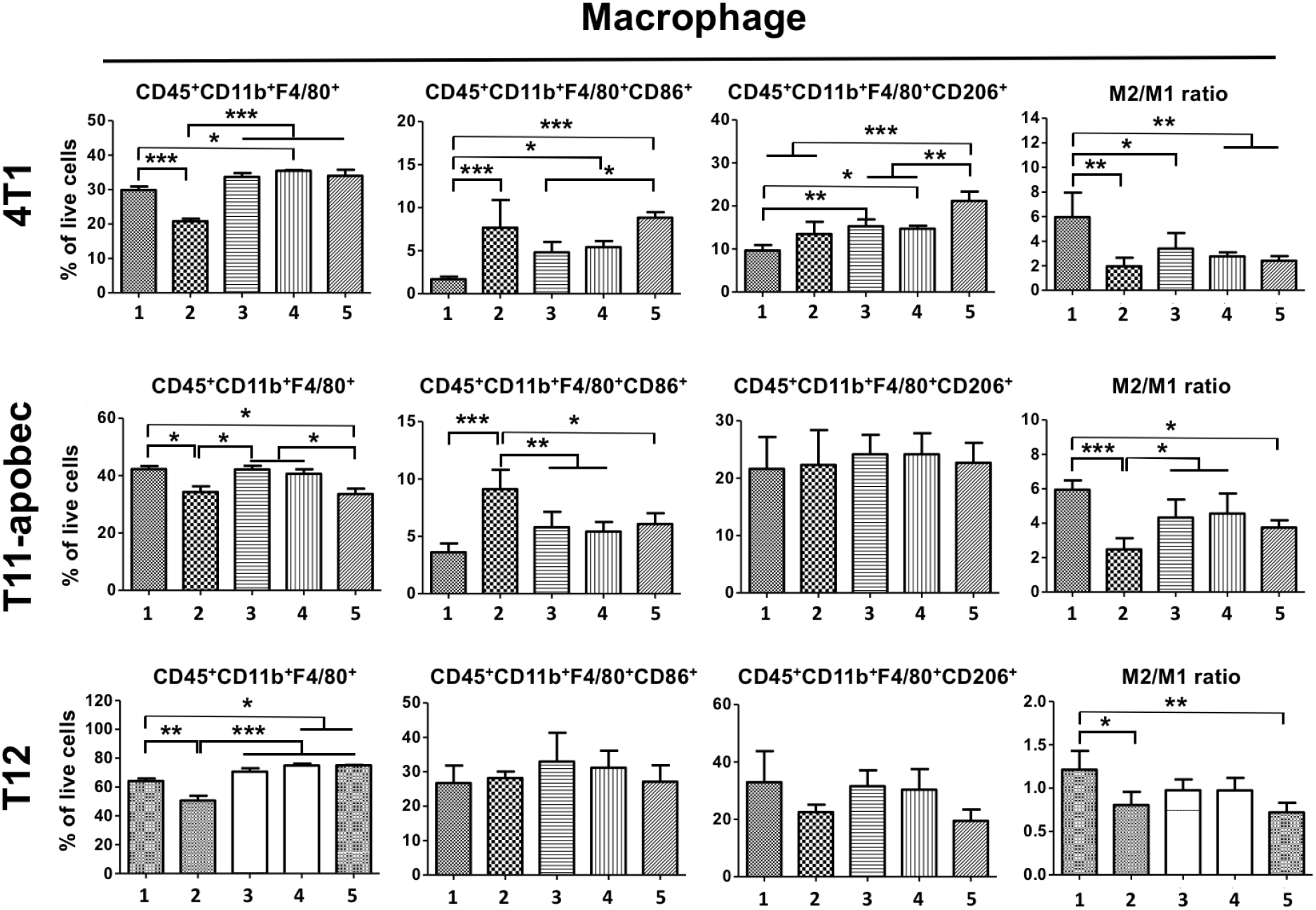
Immune phenotyping to demonstrate the effect of drug formulations on TNBC tumor models. The animals received saline or drug-loaded micelles iv using q4d × 2 with 1 – normal saline, 2 –POx-PLX (75 mg/kg), 3 – POx-PTX (30 mg/kg), 4 – POx-PTX (75 mg/kg), and 5 – co-loaded POx-PTX/PLX (75 mg/kg PTX and 75 mg/kg PLX3397). 3 Days after 2^nd^ dose of treatment, tumors were harvested to perform flow cytometry to show the impact on total macrophages (CD45^+^CD11b^+^F4/80^+^), M1-like macrophages (CD45^+^CD11b^+^F4/80^+^CD86^+^), M2-like macrophages (CD45^+^CD11b^+^F4/80^+^CD206^+^), and M2/M1 ratios. Statistical comparison of data for tumor inhibition was done using a one-way analysis of variance (ANOVA) with Tukey’s test for multiple comparison (* (p < 0.05), ** (p < 0.01), and *** (p < 0.001)). (see Supplementary **Table S6** for statistical comparisons between all groups).

### 2.6 Effect of immune cell depletion on the anti-tumor activity of drug combinations

To assess a potential role of T cell-mediated anti-tumor immunity we determined whether the depletion of the CD4^+^ and CD8^+^ T cells can interfere with the therapeutic effects of our drugs on tumor growth and lung metastases (**Figure 7** and supplementary **Figure S7**). The T cell depletions by injecting specific antibodies against CD4^+^ or CD8^+^ do not seem to influence the tumor growth curves in the saline control, and POx-PTX (30 mg/kg) or POx-PTX (75 mg/kg) treatment groups (**Figure 7(a–c)**). The effects on lung metastases in these groups were also insignificant, except for the control group where the depletion of CD8^+^ T cells increased metastatic lesions. However, in animals treated with POx-PTX/PLX the depletion of either CD4^+^ or CD8^+^ T cells significantly increased the tumor growth rates and metastatic lesions ((**Figure 7(d)**). Overall, these data indicate that the effects of POx-PTX/PLX on primary tumor growth and lung metastases are CD4^+^ and CD8^+^ cell-dependent.

**Figure 7.**
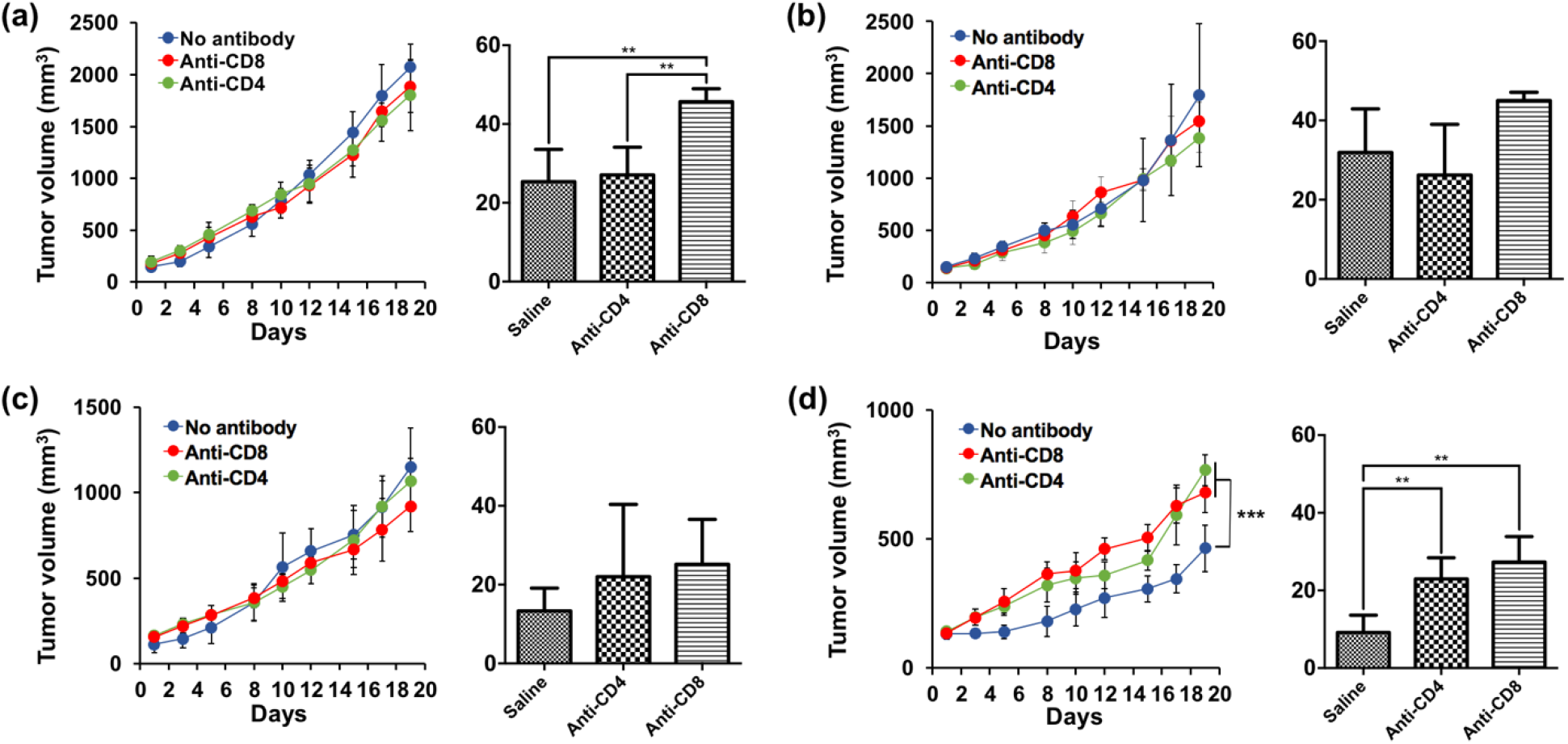
Effect of CD4^+^ and CD8^+^ cells depletion of the tumor growth curves (left panels) and lung metastasis (right panels) in 4T1 tumor bearing mice treated with: **(a)** saline, **(b)** POx-PTX (30 mg/kg). **(c)** POx-PTX (75 mg/kg) and **(d)** co-loaded POx-PTX/PLX (75 mg/kg PTX and 75 mg/kg PLX3397). Systemic depletion of CD4^+^ and CD8^+^ T cells restores tumor growth *in vivo*. Mice were treated with CD4 or CD8 depleting antibodies (intraperitoneally (ip) using q4d × 4). Two days after the first injection of an antibody, saline or drugs were administered using q4d × 4 regimen. For the assessment of the metastatic spread lungs were harvested 24 days after the first sample treatment (Supplementary Figure S8). Statistical comparisons for tumor inhibition were done using a two-way ANOVA followed by Bonferroni post-tests for multiple comparison (n = 4~5). Statistical comparisons for lung metastases were analyzed using unpaired two-tailed t-test. Statistical difference: ** (p < 0.01) and *** (p < 0.001). (see Supplementary **Table S7** for statistical comparisons between all groups)

### 2.6 Effect of the treatments of the tumor rechallenge and induction of ICD

To further demonstrate involvement of immune component in the anti-tumor responses to our drugs we carried out the vaccination-rechallenge experiment *in vivo* [49]. For this experiment, an orthotopic 4T1 tumor model was established by subcutaneously injecting 1 × 10^6^ 4T1 tumor cells into the right 4^th^ mammary fat pad. The primary tumor bearing mice received saline or drug treatments by iv injection at days −6 and −3, and the secondary 1 × 10^6^ 4T1 cells were inoculated into the left 4^th^ mammary fat pad at day 0 (**Figure 8(a)**). The secondary tumor growth effects were monitored for 14 days (**Figure 8(b)**). Notably, when the dosing amount of PTX during treatment of the primary tumor increased up to 75 mg/kg, a significant suppression of the secondary tumor growth was observed compared to saline and lower PTX dose of 30 mg/kg. The treatments using the combination drug micelles revealed a trend for further suppression of the secondary tumor although in this case the difference with the single high dose POx-PTX micelles was not significant. Based on the previously reported pharmacokinetics of POx-PTX micelles [50] at 72 h post injection the amount of drug remaining in the plasma (and tissue) is less than 0.1 μg/mL (or gram of tissue). This amount remaining after treatment of the primary tumor could not cause anti-tumor activity against the tumor challenge (IC_50_ in 4T1 is approximately 2.4 μg/mL). However, to address this point a separate tumor growth study was conducted as presented in supplementary **Figure S7**. In this experiment, the tumor-free animals were treated with two doses of the drug and then the “secondary” tumor was inoculated 3 days after the last dose of drug like the pervious experiment. There was no difference in the tumor growth rate between saline and drug treated groups. Therefore, the decrease in the secondary tumor growth observed in the vaccination-rechallenge experiment is likely due to the induction of the immunological memory because of the response elicited by the drug treatment of the primary tumor. Several studies indicated that PTX can induce ICD in TNBC and other tumors [51–53]. Therefore, we examined whether the PTX in polymeric micelles can induce ICD in 4T1 cells by measuring extracellular ATP and the surface expressions of CRT as danger-associated molecular patterns (DAMPs) associated markers [54]. As the drug dose increased the release of ATP, and expression of CRT gradually increased (**Figure S8**). To further identify the ICD, the primary tumor after second drug treatment were harvested, sectioned, and stained for CRT and HMGB1. Both markers were increased especially in the high-dose PTX and combination drug treatments suggesting that these treatments induce ICD in the tumors (**Figure 8(c)**).

**Figure 8.**
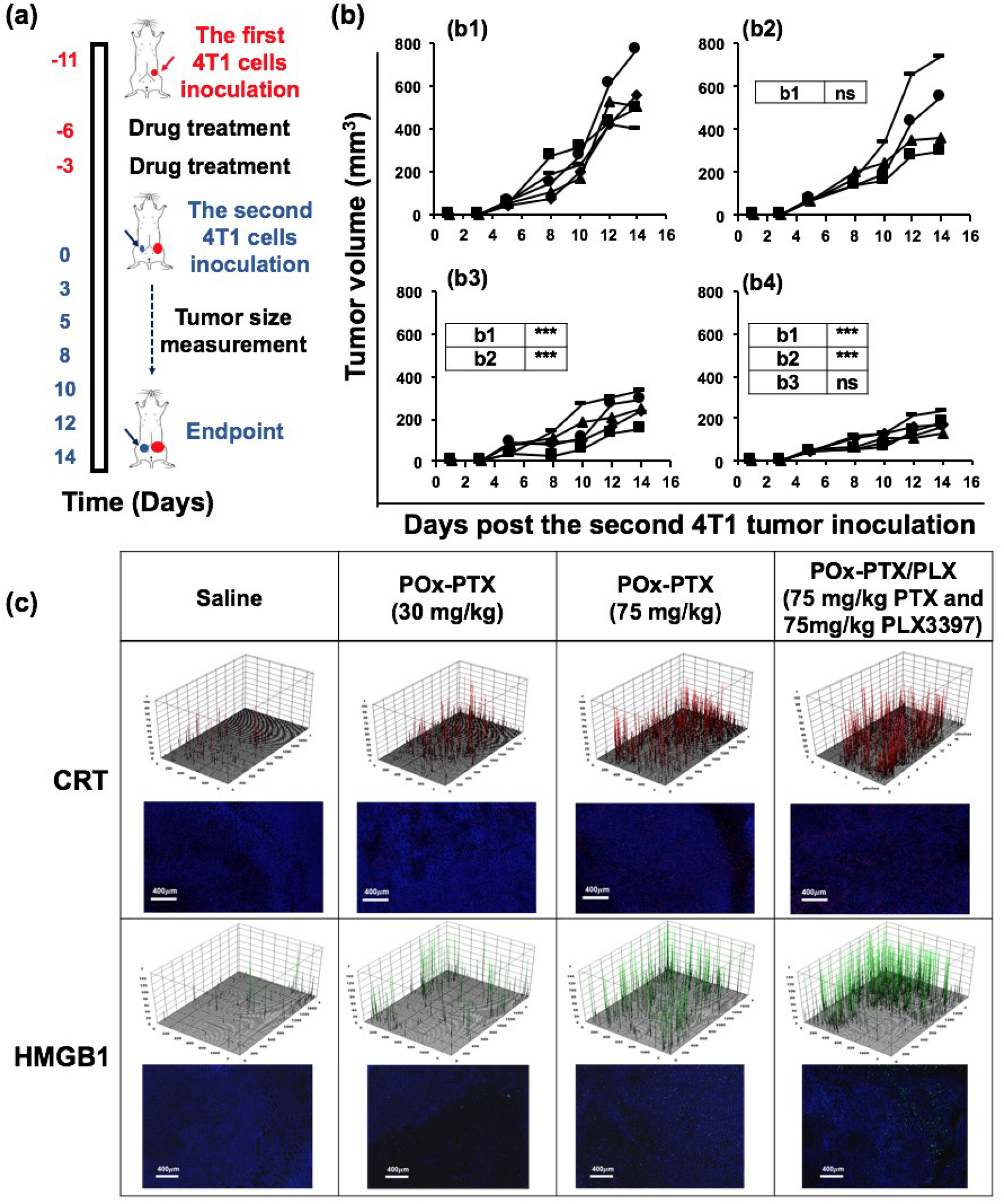
(a,b) The vaccination-rechallenge experiment and (c) evaluation of ICD *in vivo* in 4T1 breast tumor model. (a) Scheme of the vaccination protocol and (b) secondary tumor growth after treatment of the primary tumor with (b1) saline, (b2) POx-PTX (30 mg/kg), (b3) POx-PTX (75 mg/kg) and (b4) POx-PTX/PLX (75 mg/kg PTX and 75 mg/kg PLX3397). 4T1 breast cancer cells (10^6^ cells) were inoculated into 4^th^ mammary fat pad of BALB/c mice, and when the tumor sizes reached ca. 80~100 mm^3^, the animals received saline or drug-loaded micelles iv using q4d × 2 regimen. 3 Days after 2^nd^ dose of treatment, mice were rechallenged with living cancer cells of the same type, inoculated into contralateral 4^th^ mammary fat pad. The second tumor growth are routinely monitored for 14 days. Statistical difference: * (p < 0.05), ** (p < 0.01), and *** (p < 0.001). (c) Representative sections of saline or drug-loaded micelles treated tumor, immunostained for 4’,6-diamidino-2-phenylindole (DAPI), calreticulin (CRT), and high mobility group box protein 1 (HMGB1). The animals were inoculated with the primary tumor and treated as described above. Tissues were harvested 2 days after the second treatment. Stained signals were visualized by 3D surface plots in Image J.

## 3. Discussion

The use of nanotechnology approaches for the delivery of biologically active molecules has revolutionized human health as most obviously seen from the advent and implementation of RNA vaccines during the pandemic of the coronavirus infection [55, 56]. POx micelle nanoassemblies is a new platform technology that shows extremely high entrapment of small molecules with minimal amount of polymeric excipient [48]. Prior studies have shown that this technology enables to 1) deliver high-dose PTX with less toxicity than conventional paclitaxel [50], and 2) co-deliver two drugs in a single micelle with increased tumor distribution of both drugs [28, 29]. Both these effects were shown to increase outcomes of anti-cancer therapy across diverse set of cancers in rodent models.

Here, for the first time we demonstrate that high-dose chemotherapy of PTX in POx micelles potentiates the immunological effects of the drug therapy in three immune-competent mouse models of TNBC. In this study, with POx-PTX we observe enhanced tumor growth inhibition and suppression of lung metastases along with the changes in TME including increased levels of CD4^+^ and CD8^+^ T cells (in two out of three tumor models), suppression of MDSC (G-MDSC), and, notably, repolarization of TAMs toward M1-like pro-inflammatory phenotype. Moreover, POx-PTX treatment at the highest dose used (75 mg/kg) (which is still twice less than the reported MTD [50]), establishes long-term immune memory against tumor cells challenge. Notably, PTX was previously shown to induce ICD, a form of regulated cell death that triggers adaptive immunity through production of neoantigens and release of DAMPs [51–53]. PTX also can alter TAMs toward pro-inflammatory phenotype and produce anti-tumor effect by activating Toll-like receptor 4 [57]. In clinical settings the need of steroid premedication to decrease adverse effects of conventional paclitaxel may blur the anti-cancer immune responses [58]. Here, due to reformulation of PTX in POx micelles we were able to administer high-dose PTX therapy and indeed observed the dose-dependent ICD induction in the TNBC model. Notably, the greatest ICD activation in tumors as assessed by the expression of the DAMPs associated markers was seen at the highest POx-PTX dose (75 mg/kg), which is not achievable with conventional paclitaxel (MTD 20 mg/kg with q4d × 4 regimen [50]. The ICD induction along with indications of the enhanced anti-tumor immunity provides new, additional and important evidence of a potential benefit of POx micelle system that allows enhancing PTX effect on the target by delivering high dose with minimal formulation toxicity [50]. The PTX POx micelle formulation does not induce complement activation [50, 59], which may abolish in future the need of the premedication in the clinical use.

For the combination chemo- and immunotherapy, we have chosen a small molecule inhibitor of CSF-1R, PLX3397 (Pexidartinib). Pexidartinib was initially developed by Plexxicon Inc. and Daichi-Sankyo, and advanced to a multi-center international clinical trial [60]. With positive clinical outcome for the patients with symptomatic tenosynovial giant cell tumor, PLX3397 received regulatory approval in 2019 as TURALIO^®^ in United States [60]. Although this agent has been administered orally in the clinic, we assessed whether its co-formulation with PTX in polymeric micelles could be advantageous for systemic administration compared to oral dosing. Indeed, we observed improved anti-tumor effect of PTX and PLX3397 co-formulated in POx micelles and administered iv compared to the same cumulative dose of both drugs with POx-PTX injected iv and PLX3397 dosed orally in a standard vehicle. Previously, DeNardo et el. reported improved antitumor activity of iv PTX (via into the retroorbital plexus) and oral PLX3397 combination in the treatment of MMTV-PyMT TNBC [22]. However, this murine tumor model was quite sensitive to the low dose PTX therapy (10 mg/kg, q5d ×3 or q5d × 4). In contrast, all our TNBC models have shown little or no response to PTX at 30 mg/kg (q4d × 4). Across all tumor models used in this study, the co-formulated POx-PTX/PLX was more potent than the single drug treatments. Interestingly, nearly the same improved anti-tumor effect was observed with the two micellar drugs mixed before injection which may be a technological advantage for the future drug development due to ease of manufacturing and application in some clinical settings. Prior studies of chemotherapeutic agents in POx micelles, such as PTX and cisplatin prodrug derivative, suggested that these agents were more active in co-formulation [28]. Here, we observed a similar effect for the combination of chemotherapeutic and immunotherapeutic agents. Notably, PLX3397 has a direct cytotoxic effect via inhibiting c-kit and fms-like tyrosine kinase 3 pathways on cancer cells as was shown previously [33, 34]. We re-confirmed TME-independent PLX3397 cytotoxicity in our tumor cell models, and, moreover, observed strong cytotoxic synergy between PLX3397 and PTX, which provided additional rationale for combining these two drugs.

The assessment of the changes of the immune cell populations in TME, induction of ICD and tumor rechallenge experiments suggest that the co-loaded drug combination has the most robust immunotherapeutic effect compared to both single dose therapies using POx-PLX and POx-PTX. By depleting the CD4^+^ and CD8^+^ T cells we observed partial attenuation of the tumor growth suppression suggesting involvement of T cell immunity in the anti-cancer effects of POx-PTX/PLX. Moreover, we found that POx-PTX/PLX efficiently suppressed the metastatic spread of TNBC (4T1) toward the lungs. Notably, the high-dose POx-PTX has also shown significant activity in this context. This finding is quite remarkable in view of previous conflicting reports suggesting that potential involvement of the anti-tumor immune responses by PTX may promote metastases while inhibiting the primary tumor [61, 62]. The strong anti-metastatic activity of both POx-PTX and POx-PTX/PLX may be of future clinical significance for management of life-threatening metastasis in breast cancer.

In conclusion, this study adds a new perspective for chemoimmunotherapy of TNBC using POx polymeric micelles. The results demonstrate that PTX even as a single agent displays strong effects on TME and induces the long-term immune memory. We suggest that the POx micelle formulation has a potential of transforming the use of this well-known drug by enabling its high dose therapy. In addition, we demonstrate that the combination chemoimmunotherapy using PTX and PLX3397 provides consistent improvement of therapeutic outcomes across several TNBC models. These treatments are associated with the repolarization of the immunosuppressive TME and increased T cell immune responses that contribute to the suppression of both the primary tumor and metastatic disease. Overall, the work provides evidence of benefit of reformulation and outlines potential translational path for both PTX and PTX and PLX3397 combination using POx polymeric micelles for the TNBC.

## 4. Materials and methods

### 4.1. Materials

Amphiphilic triblock copolymer of poly (MeOx_35_-*b*-BuOx_20_-*b*-MeOx_34_) (M_n_ = 8.6 kDa, M_w_/M_n_ = 1.15) was synthesized by living cationic ring-opening polymerization as described previously [31, 32]. Structural properties of POx were determined by ^1^H NMR spectroscopy (INOVA 400) and gel permeation chromatography (GPCmax VE-2001 system (Viscotek)). PTX was purchased from LC laboratories (Woburn, MA). PLX3397 was purchased from MedKoo Biosciences (Morrisville, NC). All other chemicals were from Fisher Scientific INC. (Fairlawn, NJ) and of analytical grade. 4T1 cells were obtained from UNC Lineberger Tissue Culture Facility. T11-apobec and T12 cells were provided by Dr. Charles M. Perou (Lineberger Comprehensive Cancer Center, Chapel Hill, NC, USA). 4T1 cells were cultured in RPMI medium (11965-092 (Gibco)) supplemented with 10% fetal bovine serum (FBS) and 1% penicillin-streptomycin at 37 °C in a cell culture incubator. T11-apobec cells were cultured in RPMI medium (11965-092 (Gibco)) supplemented with 5% FBS, 1% penicillin-streptomycin and puromycin. T12 cells cultured in RPMI medium (11965-092 (Gibco)) supplemented with 5% FBS and 1% penicillin-streptomycin.

### 4.2. Preparation and characterization of drug-loaded polymeric micelle formulations

POx micelle formulations with drugs were prepared by thin-film hydration method as previously described [28, 29, 31, 50]. Briefly, stock solutions of the POx polymer and drugs (PTX and PLX3397) were prepared in absolute ethanol (at 10 mg/mL POx, 5 mg/mL PTX and 2 mg/mL PLX3397, respectively). These stock solutions were thoroughly mixed at the pre-determined ratios based on drug feeding ratio in POx micelles. By completely evaporating of ethanol under a stream of inert nitrogen gas, thin films of drug-polymer homogeneous mixture were obtained. Thin films were subsequently hydrated with sterile saline and incubated (at 70 °C for POx-PTX (10 min) and POx-PTX/PLX (5 min), and at room temperature for POx-PLX (10 min)). The resulting micelle formulations were centrifuged at 10,000 g for 3 minutes (Sorvall Legend Micro 21R Centrifuge, Thermo Scientific) to remove any drug precipitates from the micelle formulations. Clear supernatants of the micelle solutions were collected and used for physicochemical analysis of the micelle drugs.

Drug loadings in given micelle drugs were analyzed by high-pressure liquid chromatography (HPLC) system (Agilent Technologies 1200 series) using a Nucleosil C18 column (4.6 mm × 250 mm, 5 μm). A mobile phase composed of water (0.1% trifluoroacetic acid) and acetonitrile (0.1% trifluoroacetic acid) (50/50 volume ratio) was used. Micelle formulations were diluted 50-fold with mobile phase and 10 μL of diluted micelle samples were injected to HPLC system for drug loading analysis. Flow rate of the mobile phase was 1.0 mL/min. Detection wavelength were 270 nm for PLX3397 and 235 nm for PTX. The LE and LC of the micelle formulations were calculated using following equations (1)–(2).

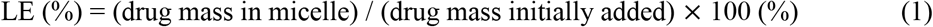

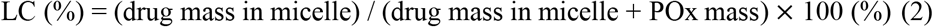

Size distribution of micelle drugs in solution was measured by Zetasizer Nano ZS (Malvern Instruments Ltd., UK) equipped with a multi-angle sizing option. Effective diameter (Deff) and PDI of each micelle drugs were determined by DLS from three measures of three independently prepared micelle drugs. The morphology of POx-PTX/PLX was examined using a LEO EM910 TEM operating at 80 kV (Carl Zeiss SMT Inc., Peabody, MA). Dilute samples of POx-PTX/PLX were attached to copper grid/carbon film and stained with 1% uranyl acetate prior to the TEM imaging. Digital images were obtained using a Gatan Orius SC1000 CCD Digital Camera in combination with Digital Micrograph 3.11.0 software (Gatan Inc., Pleasanton, CA).

The drug release from micelle drugs were investigated using the membrane dialysis method under sink condition [28, 29, 31, 50]. Briefly, micelle drugs were diluted in phosphate-buffered saline (PBS) to achieve 0.1 g/L of total drug concentration in the solution. Then the diluted micelle drug solutions were transferred to floatable Slide-A-Lyzer MINI dialysis devices (100 μL capacity, 20 kDa MWCO) and placed in 30 mL of PBS supplemented with 10% FBS. At predetermined time points, four devices were sacrificed and micelle drug solutions in the devices were collected. Remaining drugs in the obtained micelle drug samples were quantified by HPLC as described above. The drug release profiles were plotted by expressing % drug released from micelle drug over time.

### 4.3. *In vitro* cytotoxicity and induction of immunogenic cell death

*In vitro* cytotoxicity by micelle drugs was determined by the measurement of cell viability using CCK-8 assay (Dojindo, MD). A 96-well cell culture plates were used to seed 4T1, T11-Apobec, and T12 cells (1 × 10^4^ cells/well) for 24 h at 37 °C in an atmosphere of 5% CO2 with fresh media. Following 24 h incubation of the TNBC cells in the plates, the cells were treated with micelle drugs of serial dilutions for 48 h. The cells in the plates were then washed with sterile PBS, CCK-8 agents were added to the plates, and the plates were incubated at 37°C in an atmosphere of 5% CO2 for 4 h. The absorbance at 450 nm of each wells of the plates was recorded on a UV spectrophotometer (SpectraMax M5, Molecular Devices). Cytotoxicity of micelle drugs on the TNBC cells (expressed as IC_50_) were analyzed using the Graphpad Prism 7.03 software.

The dead cell apoptosis kit with Annexin V (V13242 (Invitrogen)) was used to analyze the apoptosis and necrosis [63] induced by micelle drugs. 4T1 cells (5 × 10^4^ cells/well) were seeded in 24-well plates and treated with micelle drugs at 1 μg/mL of total drug concentration. After 24 h, the 4T1 cells were harvested and double-stained with Annexin V-FITC and propidium iodide as mentioned in the product manuals. The apoptotic/necrotic 4T1 cells were analyzed by Attune NxT flow cytometer (Thermo Fisher) and the portions of apoptotic/necrotic 4T1 cells were analyzed using FlowJo v10.4.2 software.

To investigate the induction of ICD *in vitro* by micelle drug treatment, ICD markers such as ATP and CRT measured upon treatment of micelle drugs on 4T1 cells. For ATP concentration measurement *in vitro*, 4T1 cells (1 × 10^5^) were seeded in 24-well plates in fresh media for 24 h, then treated with micelle drugs with serial dilutions for 24 h. Media supernatants of the 4T1 cells in the plates were collected and immediately analyzed with ATPlite one-step luminescence assay kits (PerkinElmer, MA) for the measurement of ATP concentration in the media. For the detection of CRT translocation to the cell surface by ICD [49], 4T1 cells (1 × 10^5^) were seeded in 24-well plates in fresh media for 24, then treated with micelle drugs with serial dilutions for 24 h. Subsequently, the 4T1 cells in the plates were fixed with 4% formaldehyde and stained with anti-CRT antibody (ab196159 (Abcam)). CRT^+^ 4T1 cells were cells were analyzed by Attune NxT flow cytometer (Thermo Fisher) and the percentage of CRT^+^ 4T1 cells were analyzed using FlowJo v10.4.2 software.

### 4.4. Determination of MTD

Animal studies were conducted in accordance with the University of North Carolina at Chapel Hill Institutional Animal Care and Use Committee guidelines. MTD of micelle drug (POx-PTX/PLX) was determined by dose-escalation study using healthy 8-weeks old female BALB/c mice (Jackson Laboratory). Mice were randomly divided into five groups and POx-PTX/PLX was administered iv using either q4d × 4 or q4d × 6 regimens. The mice were injected iv with: saline, POx-PTX/PLX (75 mg/kg PTX and 75 mg/kg PLX3397) (iv; q4d × 4), POx-PTX/PLX (100 mg/kg PTX and 100 mg/kg PLX3397) (iv; q4d × 4), POx-PTX/PLX (125 mg/kg PTX and 125 mg/kg PLX3397) (iv; q4d × 4), or POx-PTX/PLX (75 mg/kg PTX and 75 mg/kg PLX3397) (iv; q4d × 6). The body weight change of the mice was monitored every day for 27 days. Drug treatments were discontinued if any signs of toxicity behavior including hunched posture, rough coat and body weight changes over 15% of the initial body weight.

### 4.5. *In vivo* TNBC animal models and tumor inhibition studies

Animal studies were conducted in accordance with the University of North Carolina at Chapel Hill Institutional Animal Care and Use Committee guidelines. In order to investigate tumor inhibition by micelle drug treatments, three orthotopic TNBC models (4T1, T11-apobec, and T12) were employed. Tumor-bearing mouse models were developed as follows. For 4T1 model, 6-8 weeks old female BALB/c mice (Jackson Laboratory) were orthotopically inoculated in the 4^th^ mammary fat pad with 4T1 cells (1 x 10^6^ cells in 100 μL of PBS and Matrigel (Corning, AZ) mixture (1:1 volume ratio)). For T11-apobec model, female BALB/c mice (6-8 weeks) were orthotopically inoculated in the 4^th^ mammary fat pad with T11-apobec cells (1 x 10^5^ cells in 100 μL of PBS and Matrigel mixture (1:1 volume ratio)). For T12 model, female BALB/c mice (6-8 weeks) were orthotopically inoculated in the 4^th^ mammary fat pad with T12 cells (1 x 10^5^ cells in 100 μL of PBS and Matrigel mixture (1:1 volume ratio)). For all breast cancer models, when the tumor sizes reached ca. 100 mm^3^, animals were randomized (n = 5) and received the following iv injections via tail vein using q4d × 4 regimen. 1) sterile saline; 2) POx-PTX (30 mg/kg PTX); 3) POx-PLX (75 mg/kg PLX3397); 4) POx-PTX (75 mg/kg PTX); 5) POx-PTX/PLX (30 mg/kg PTX and 30 mg/kg PLX3397), and 6) POx-PTX/PLX (75 mg/kg PTX and 75 mg/kg PLX3397). Tumor size was closely monitored every 2–3 days and tumor length (L) and width (W) were measured, and tumor volume (V) was calculated using the following equation: V = ½ x L x W^2^. 4T1 tumor bearing-mice were sacrificed on day 21, and tumor samples and lung samples were collected for IHC analysis. For oral gavage, PLX3397 was solubilized in the mixture of 5% DMSO, 45% PEG300, and 5% Tween 80 in distilled water. For iv administration of free drugs, PTX and PLX3397 were solubilized in the mixture of 50% ethanol and 50% Cremophor, and diluted 5 times in PBS before use.

For the CD4^+^ and CD8^+^ T cell depletion, 4T1 tumor-bearing BABL/c mice were treated with ip injections of an anti-CD4 or anti-CD8 antibody or control saline 2 days before each drug treatment for 4 times in total.

### 4.6. Lung metastasis quantification

Brightfield whole slide images of hematoxylin and eosin-stained lung sections were scanned at 20-fold magnification by the UNC Pathology Services Core (PSC) using the Aperio AT2 digital scanner (Leica Biosystems Imaging, Inc., Deer Park, IL). Digital image analysis was performed by a boarded veterinary pathologist in the UNC PSC using Definiens Architect XD 64 version 2.7.0.60765 to detect tumors within the lung parenchyma. First, lung (ROIs) was detected automatically with a minimum tissue size of 150,000 *μ*m^2^, 221 brightness threshold, and 3.6 homogeneity threshold. Large vessels, bronchi, and thymus tissue were manually excluded from the initial ROI. Lung components were segmented into lung, glass (alveolar spaces), blood (erythrocytes in large vessels or alveolar spaces), tumor, and outside lung (glass on the outside of the lung) categories. The resulting lung and blood area (*μ*m^2^) for each whole slide image were added to calculate the total tissue area. The tumor area (*μ*m^2^) was then divided by the total tissue area to obtain the tumor percent area. The algorithm was validated by a semi-quantitative assessment of relative % tumor burden for a subset of the slides.

### 4.7. Flow cytometry

For analyzing the immune cell population changes in the tumors (4T1, T11-apobec, and T12) upon micelle drug treatments, orthotopic primary tumors were harvested 3 days after the second dose. The harvested tumors were enzymatically digested with collagenase (2 mg/ml in Hank’s balanced salt solution (HBSS)), dispase (2.5 U/ml in HBSS), and deoxyribonuclease (1 mg/ml in PBS) for 45 min while shaking in incubator at 37°C, and then passed through a 40 *μ*m cell strainer.

The cells were treated with ammonium-chloride-potassium buffer to lyse the red blood cells and dispersed in FACS buffer (2 % FBS in PBS solution). Single cell suspensions were counted and live cells (1 × 10^6^) were stained with a cocktail of fluorescently labeled antibodies for the pre-designed panel (see supplementary **Table S4** and **S5)**. Antibodies panel used for flow cytometry are listed in supplementary **Table S5**. Then the cells were fixed with 4% paraformaldehyde solution and analyzed with Attune NxT flow cytometer (Thermo Fisher, MA) at UNC Flow Cytometry Core Facility. Cell population analysis was performed using FlowJo software (TreeStar, Ashland, OR).

### 4.8. *In vivo* immunogenic cell death vaccination study

Schematic presentation of the vaccination protocol and drug treatment regimen for the vaccination-rechallenge experiments is described in **Figure 8(a)**. Briefly, female BALB/c mice (6-8 weeks old) were orthotopically inoculated with 4T1 cells (1 × 10^6^ cells in 100 μL of PBS and Matrigel mixture (1:1 volume ratio)) in the 4^th^ mammary fat pad. When the tumor sizes reached ca. 50~80 mm^3^, the animals received the following micelle drugs via iv injections using q4d × 2 regimen. 1) saline; 2) POx-PTX (30 mg/kg PTX); 3) POx-PTX (75 mg/kg PTX); 4) POx-PTX/PLX (75 mg/kg PTX and 75 mg/kg PLX3397). Three days after 2^nd^ dose of treatment, mice were rechallenged with living 4T1 cancer cells of the same type, inoculated into contralateral 4^th^ mammary fat pad. The second tumor growths were routinely monitored for 14 days. Secondary tumor volume (V) was calculated using the following equation: V = ½ x L × W^2^. Mice were sacrificed on day 14 from the secondary tumor inoculation, and the tumors were collected for IHC analysis. The vaccination rechallenge experiments were also conducted with healthy BALB/c mice (see schematic presentation of the protocol in **Figure S7(a)**). Briefly, Healthy mice were treated with saline or POx-PTX/PLX iv using q4d × 2 regimen. 3 days after 2nd dose of treatment, mice were challenged with living cancer cells of 4T1 TNBC, inoculated into 4th MFP. The primary tumor growth was routinely monitored for 14 days.

### 4.9. Statistical analysis

GraphPad 6.0 was used for statistical analysis in this study and numerical results are expressed as mean ± standard deviation in figures and tables. For comparison within two groups, unpaired two-tailed t-test was used. For comparison among multiple groups, one-way analysis of variance (one-way ANOVA) or two-way ANOVA by Bonferroni post-tests were employed for statistical analysis. Animal survival is presented as Kaplan-Meier survival curves and analyzed via Log-rank (Mantel-Cox) test. Following symbols are presented in the figures to represent the statistical significance: * (p < 0.05), ** (p < 0.01), *** (p < 0.001), and **** (p < 0.0001), ns (not significant).

## Acknowledgements

This work was partially supported by the National Cancer Institute (NCI) Alliance for Nanotechnology in Cancer (U54CA198999, Carolina Center of Cancer Nanotechnology Excellence), and NCI’s grant R01CA264488 (to AVK). Partial support was also provided by NCI Breast SPORE program (P50-CA058223), and RO1-CA148761 (to CMP). Animal Studies were performed within the UNC Lineberger Animal Studies Core (ASC) Facility at the University of North Carolina at Chapel Hill. YM and HH postdoctoral fellowships were supported through the Carolina Cancer Nanotechnology Training Program funded by NCI (T32CA196589). The UNC Lineberger Animal Studies Core was supported, in part, by an NCI Center Core Support Grant (CA16086) to the UNC Lineberger Comprehensive Cancer Center. The authors thank C. Santos, M. Ross, and A. Valdivia at Animal Studies Core of UNC for helping with the intravenous/intraperitoneal injections. TEM was performed by A. Shankar Kumbhar at the Chapel Hill Analytical and Nanofabrication Laboratory (CHANL), a member of the North Carolina Research Triangle Nanotechnology Network (RTNN), which was supported by the NSF (grant ECCS-1542015) as part of the National Nanotechnology Coordinated Infrastructure (NNCI).

