## Supplementary Materials for "High-Dose Paclitaxel and its Combination with CSF1R Inhibitor in Polymeric Micelles for Chemoimmunotherapy of Triple Negative Breast Cancer"

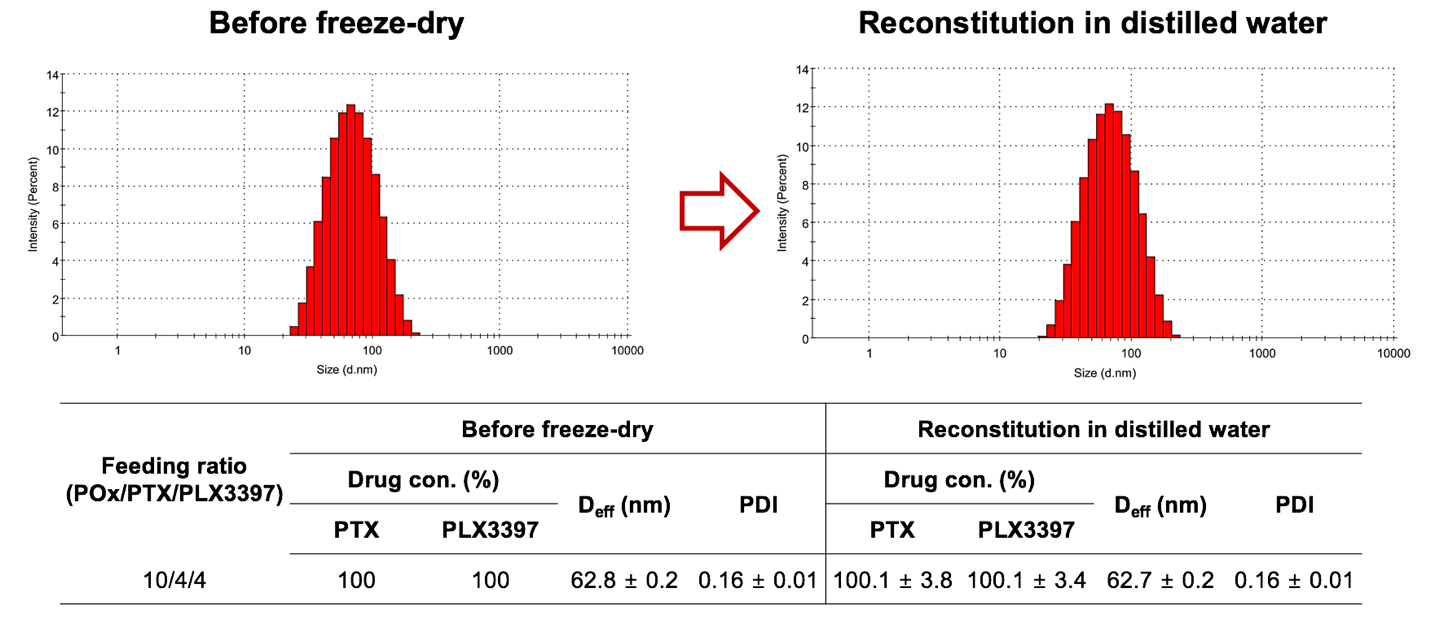

**Figure S1. Reconstitution of freeze-dried POx-PTX/PLX with distilled water and comparison of size distribution of the micelle drug before and after the reconstitution by DLS.**

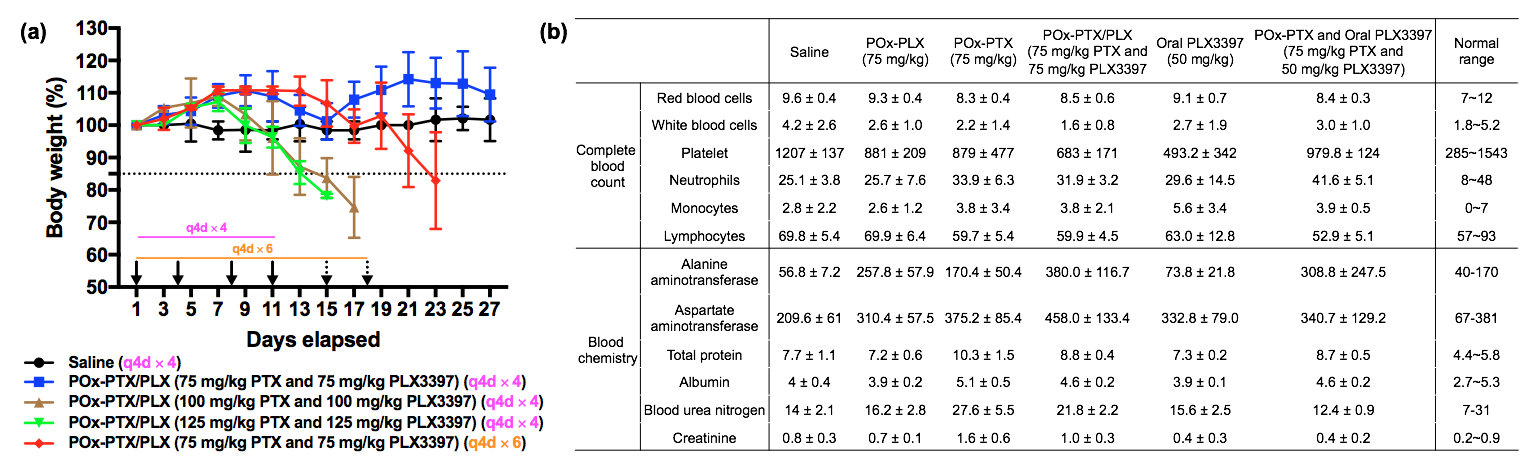

**Figure S2. PTX and PLX3397 toxicity in healthy mice.** (a) The body weight change of healthy mice injected iv (as shown by arrows) with: saline or co-loaded POx-PTX/PLX drugs as indicated dose and frequency of each drug over time. (b) Complete blood count and clinical chemistry parameters of healthy mice treated with the following formulations: normal saline (iv; q4d $\times$ 4), POx-PLX (75 mg/kg) (iv; q4d $\times$ 4), POx-PTX (75 mg/kg) (iv; q4d $\times$ 4), POx-PTX/PLX (75 mg/kg PTX and 75 mg/kg PLX3397) (iv; q4d $\times$ 4), Oral PLX3397 (50 mg/kg) (oral; q2d $\times$ 6) (dissolved in 5% DMSO, 45% PEG300, and 5% Tween 80 in distilled water), or POx-PTX and Oral PLX3397 (POx-PTX (75 mg/kg) (iv; q4d $\times$ 4) and oral PLX3397 (50 mg/kg) (oral; q2d $\times$ 6) (dissolved in 5% DMSO, 45% PEG300, and 5% Tween 80 in distilled water)).

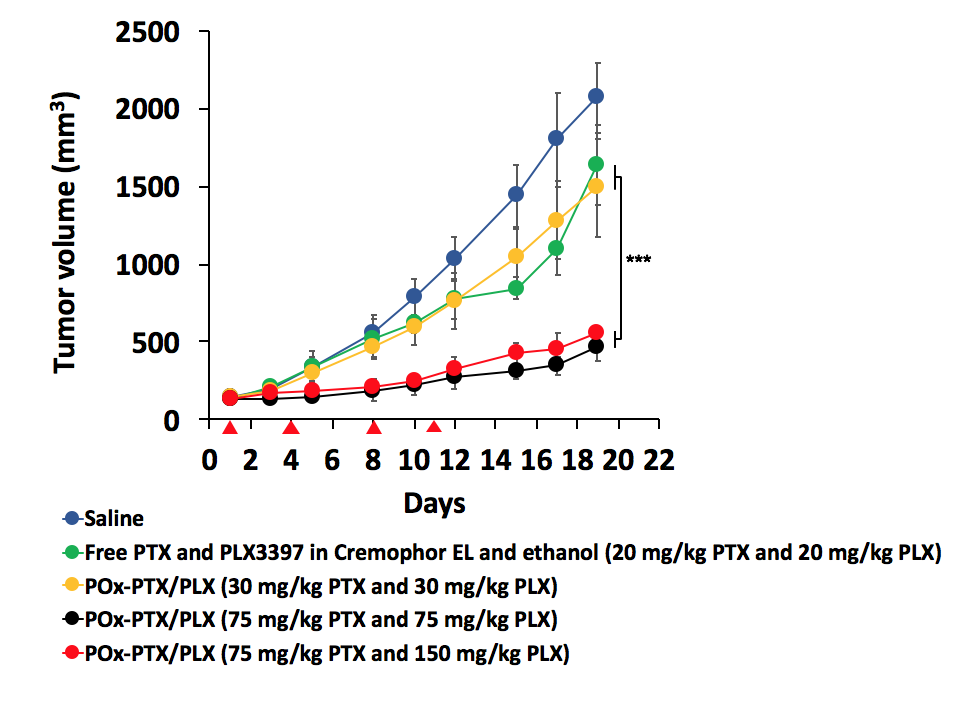

**Figure S3. Primary tumor inhibition in mice bearing orthotopic TNBC 4T1 tumor.** The animals (n = 4~8) were injected iv (as shown by arrows, q4d x 4) with: saline, free PTX and PLX3397 in Cremophor EL and ethanol (20 mg/kg PTX and 20 mg/kg PLX3397), POx-PTX/PLX (30 mg/kg PTX and 30 mg/kg PLX), POx-PTX/PLX (75 mg/kg PTX and 75 mg/kg PLX), and POx-PTX/PLX (75 mg/kg PTX and 150 mg/kg PLX). Statistical comparison of data for tumor inhibition was done using a two-way analysis of variance (ANOVA) and followed by Bonferroni post-tests for multiple comparison (n=4~5). Statistical difference: *** (p < 0.001). See **Supplementary Table S2** for the complete statistical comparison between all groups.

Some slowdown of the tumor growth compared to the saline control was observed in the groups treated with either the free drugs combination (20 mg/kg PTX and 20 mg/kg PLX) or POx-PTX/PLX at a lower dose (30 mg/kg PTX and 30 mg/kg PLX). Although this effect was significant (see supplementary **Table S2**) it was much less than that of the POx-PTX/PLX (75 mg/kg PTX and 75 mg/kg PLX) treatments. Further increase in the PLX3397 dose in the co-loaded polymeric micelle formulation within the limits that were still safe to the animal (POx-PTX/PLX (75 mg/kg PTX and 150 mg/kg PLX)), did not increase the antitumor effect in any measurable way.

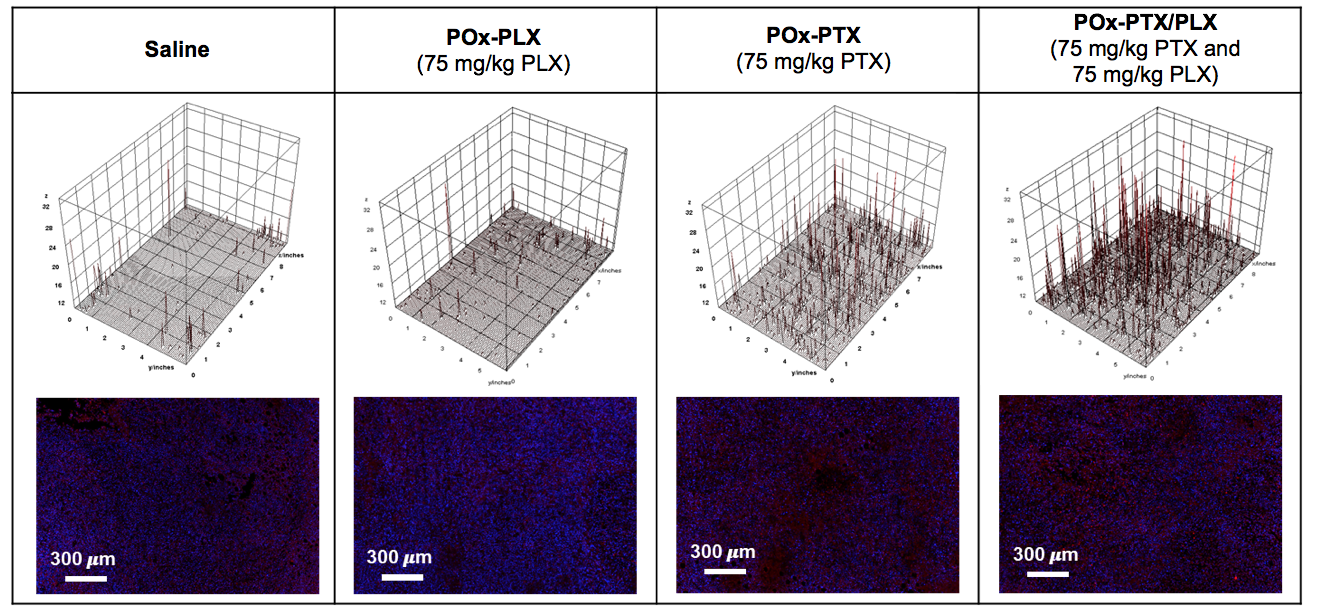

**Figure S4. Immunohistochemistry staining of tumor apoptosis.** Representative sections of saline, or drug-loaded micelles treated tumor, immunostained for DAPI, and cleaved caspase 3 (Red). The animals were inoculated with the primary tumor and treated with: saline, POx-PLX (75 mg/kg PLX), POx-PTX (75 mg/kg) and POx-PTX/PLX (75 mg/kg PTX and 75 mg/kg PLX). The tumor-bearing animals were treated with the drugs twice and tumors samples were harvested 2 days after the last second dose. Stained signal for cleaved caspase 3 were visualized by 3D surface plots in Image J.

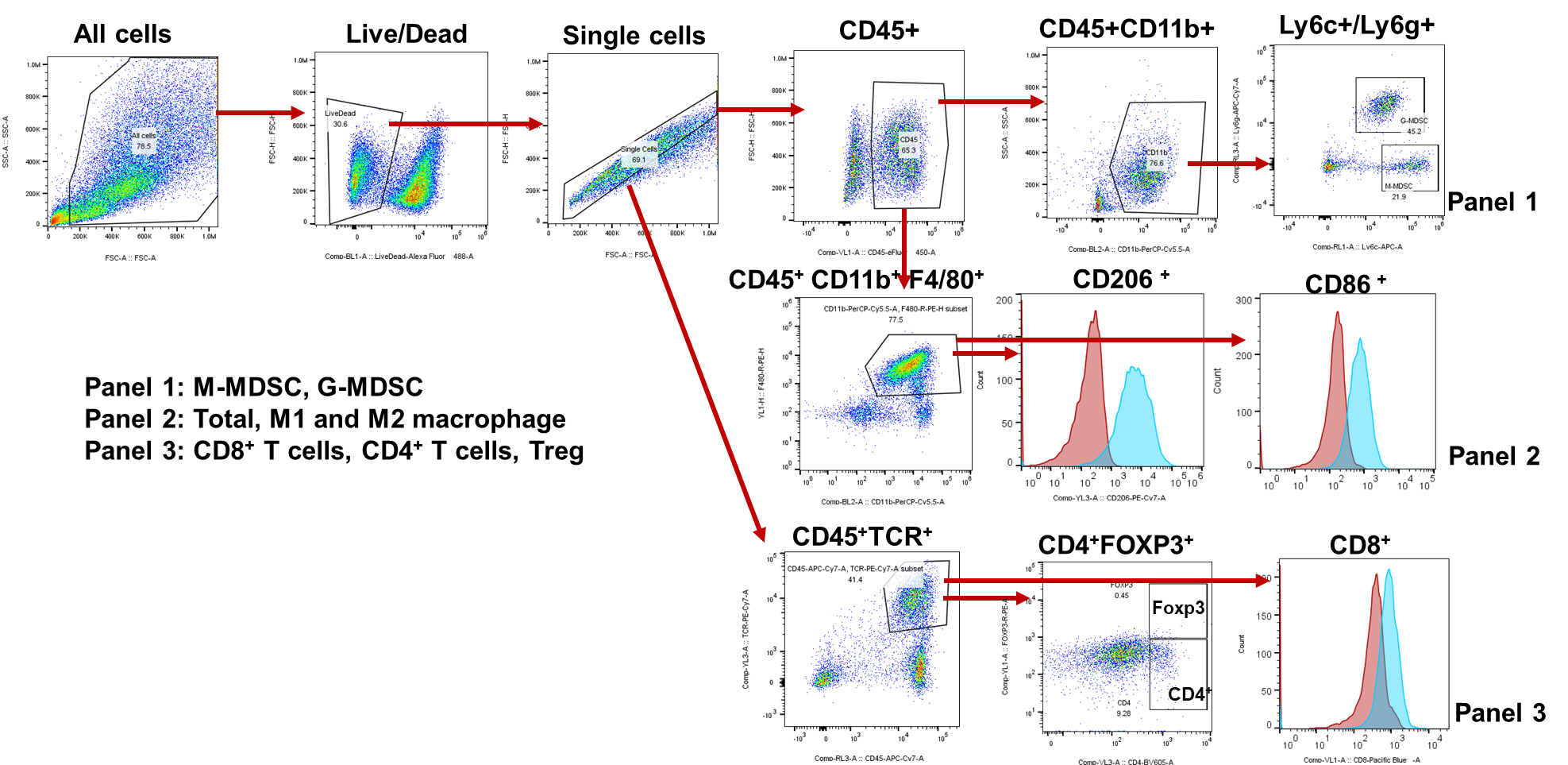

**Figure S5.** Gating strategy for multi-panel parameter flow cytometry. Fluorescence Minus One (FMO) controls for all the fluorophores in the panels were used to properly interpret flow cytometry data.

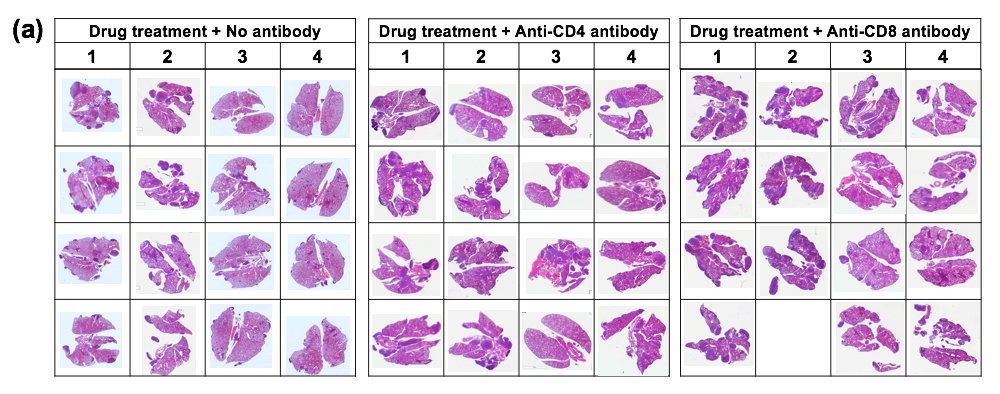

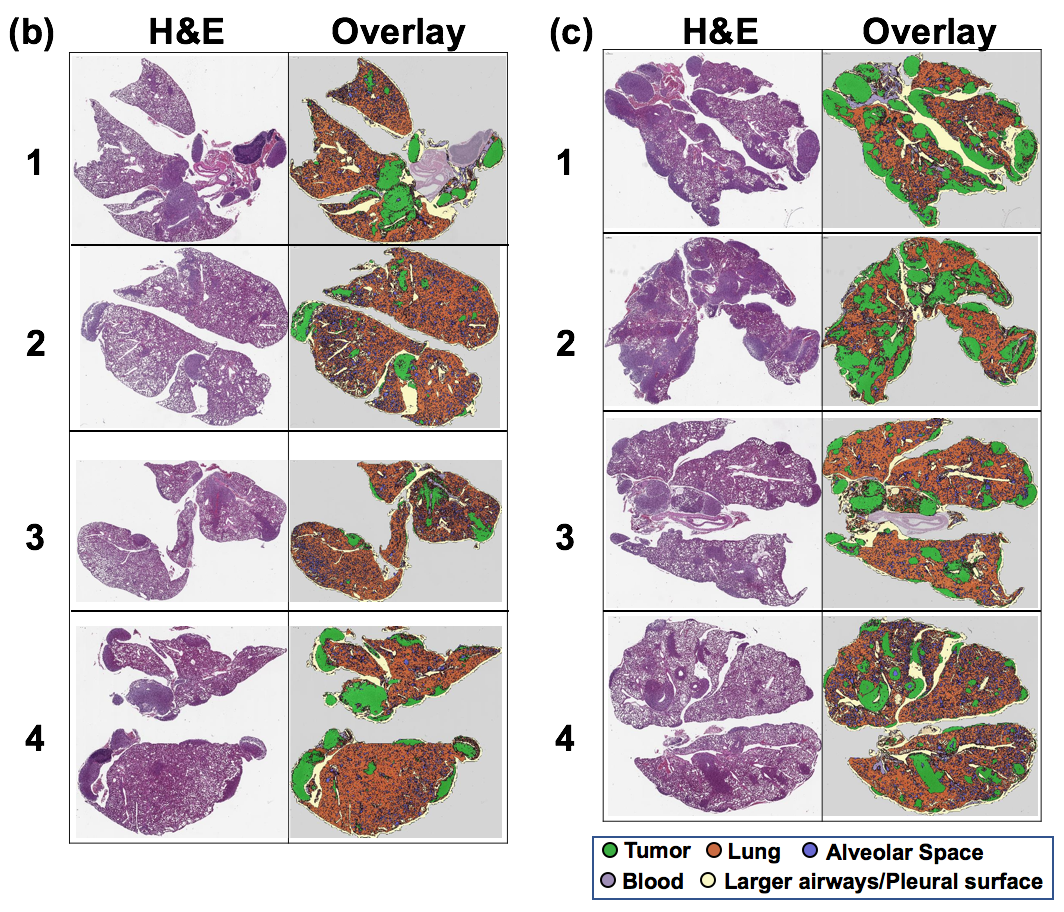

**Figure S6. Images of sections of lungs harvested from orthotopic TNBC 4T1-bearing mouse.** 4T1-bearing mice were grouped, according to antibody treatment, into three groups (no antibody, Anti-CD4, or Anti-CD8). Each group were further subdivided by drug treatments as follows: 1 – normal saline, 2 – POx-PTX (30 mg/kg), 3 – POx-PTX (75 mg/kg), 4 – co-loaded POx-PTX/PLX (75 mg/kg PTX and 75 mg/kg PLX3397) (n = 3~5). (a) H&E images of lungs in all groups, and (b) representative H&E images and overlay images of lung section with the identification of metastatic spread of 4T1 tumor in (b) Anti-CD4 group and (c) Anti-CD8 group.

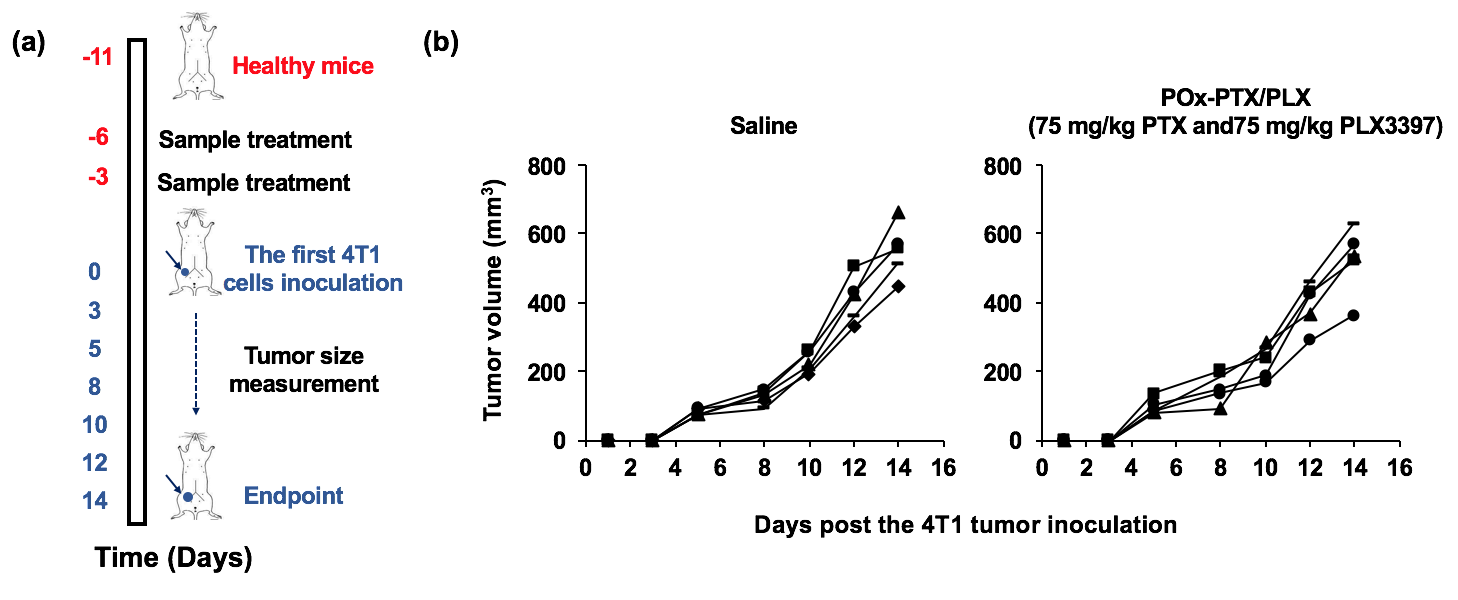

**Figure S7. The vaccination-rechallenge experiments.** (a) Scheme of the vaccination protocol and (b) primary tumor growth after treatment of healthy mice with saline, or POx-PTX/PLX (75 mg/kg PTX and 75 mg/kg PLX3397). Healthy mice were treated with saline or POx-PTX/PLX iv using q4d × 2 regimen. 3 days after 2^nd^ dose of treatment, mice were challenged with living cancer cells of 4T1 TNBC, inoculated into 4^th^ MFP. The primary tumor growths are routinely monitored for 14 days.

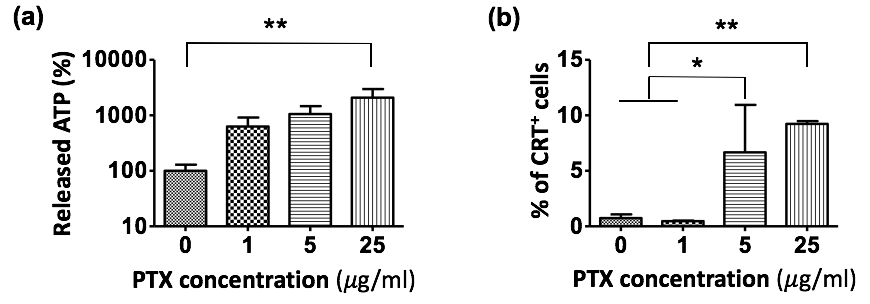

**Figure S8. Evaluation of ICD *in vitro*.** (a) Extracellular ATP level by a luciferase-based assay and (b) the assessment of CRT exposure (expressed as % of CRT^+^ cells) on the surface of live cells by flow cytometry. Statistical comparison was done using a one-way analysis of variance (ANOVA) and followed by Bonferroni post-tests for multiple comparison (n = 4~5). Statistical difference: * (p < 0.05), ** (p < 0.01).

**Table S1.** **Statistical comparisons for primary TNBC tumor inhibition between cohorts in Figure 3(a-c).** (By one-way ANOVA with Tukey’s test for multiple comparisons. Statistical difference: * (p < 0.05), ** (p < 0.01), and *** (p < 0.001). (Graphpad Prism, version 7.03).

|  | | POx-PLX  (75 mg/kg) | | POx-PTX  (30 mg/kg) | | POx-PTX  (75 mg/kg) | POx-PTX/PLX (30 mg/kg PTX  and 30 mg/kg PLX) | | POx-PTX/PLX (75 mg/kg PTX  and 75 mg/kg PLX) |
| --- | --- | --- | --- | --- | --- | --- | --- | --- | --- |
| 4T1 | Saline | ns | | ns | | *** | *** | | *** |
|  | POx-PLX (75 mg/kg) |  | | *** | | *** | *** | | *** |
|  | POx-PTX (30 mg/kg) |  | |  | | *** | ns | | *** |
|  | POx-PTX (75 mg/kg) |  | |  | |  | * | | *** |
|  | POx-PTX/PLX (30 mg/kg PTX and 30 mg/kg PLX) |  | |  | |  |  | | *** |
| T11-apobec | Saline | *** | | *** | | *** | *** | | *** |
|  | POx-PLX (75 mg/kg) |  | | ns | | ns | ns | | *** |
|  | POx-PTX (30 mg/kg) |  | |  | | ns | ns | | *** |
|  | POx-PTX (75 mg/kg) |  | |  | |  | ns | | *** |
|  | POx-PTX/PLX (30 mg/kg PTX and 30 mg/kg PLX) |  | |  | |  |  | | *** |
| T12 | Saline | *** | | ** | | *** | *** | | *** |
|  | POx-PLX (75 mg/kg) |  | | ns | | ns | ns | | *** |
|  | POx-PTX (30 mg/kg) |  | |  | | *** | ** | | *** |
|  | POx-PTX (75 mg/kg) |  |  | | |  | ns | | ns |
|  | POx-PTX/PLX (30 mg/kg PTX and 30 mg/kg PLX) |  |  | |  | | |  | ** |

**Table S2**. **Statistical comparisons primary for TNBC tumor inhibition between cohorts in Figure S3.** (By one-way ANOVA with Tukey’s test for multiple comparisons. Statistical difference: * (p < 0.05), ** (p < 0.01), and *** (p < 0.001). (Graphpad Prism, version 7.03).

|  | PTX (20 mg/kg) and PLX3397 (20 mg/kg) in Cremophor EL/ethanol | POx-PTX/PLX (30 mg/kg PTX and 30 mg/kg PLX) | POx-PTX/PLX (75 mg/kg PTX and 75 mg/kg PLX) | POx-PTX/PLX (75 mg/kg PTX and 150 mg/kg PLX) |
| --- | --- | --- | --- | --- |
| Saline | *** | *** | *** | *** |
| PTX (20 mg/kg) and PLX3397 (20 mg/kg) in Cremophor EL/ethanol |  | ns | *** | *** |
| POx-PTX/PLX (30 mg/kg PTX  and 30 mg/kg PLX) |  |  | *** | *** |
| POx-PTX/PLX (75 mg/kg PTX  and 75 mg/kg PLX) |  |  |  | ns |

**Table S3.** **Statistical comparisons primary for TNBC tumor inhibition between cohorts in Figure 4.** (By one-way ANOVA with Tukey’s test for multiple comparisons. Statistical difference: * (p < 0.05), ** (p < 0.01), and *** (p < 0.001). (Graphpad Prism, version 7.03).

|  | Sequential POx-PTX (75 mg/kg) and POx-PLX (75 mg/kg) | POx-PTX (75 mg/kg) and oral PLX (50 mg/kg) | Simultaneous POx-PTX (75 mg/kg) and POx-PLX (75 mg/kg) | POx-PTX/PLX (75 mg/kg PTX and 75 mg/kg PLX) |
| --- | --- | --- | --- | --- |
| POx-PTX (75 mg/kg) | *** | *** | *** | *** |
| Sequential POx-PTX (75 mg/kg) and POx-PLX (75 mg/kg) |  | ns | *** | *** |
| POx-PTX (75 mg/kg) and  oral PLX (50 mg/kg) |  |  | *** | *** |
| Simultaneous POx-PTX (75 mg/kg) and POx-PLX (75 mg/kg) |  |  |  | ns |

**Table S4. Multi-panel parameter flow cytometry**

|  | **Target cells** | **Antibody panel** |
| --- | --- | --- |
| T cells | Cytotoxic CD8^+^ T cells (CD8^+^ T cells) | CD45^+^TCR^+^CD8^+^ |
|  | T helper cells (CD4^+^ T cells) | CD45^+^TCR^+^CD4^+^ |
|  | Regulatory T cells (Treg) | CD45^+^TCR^+^CD4^+^Foxp3^+^ |
| MDSC | Monocytic Myeloid-derived suppressor cells (M-MDSC) | CD45^+^CD11b^+^Ly6C^+^ |
|  | Granulocytic Myeloid-derived suppressor cells (G-MDSC) | CD45^+^CD11b^+^Ly6G^+^ |
| Macrophage | Total macrophage | CD45^+^CD11b^+^F480^+^ |
|  | M1 Macrophage | CD45^+^CD11b^+^F480^+^ CD86^+^ |
|  | M2 Macrophage | CD45^+^CD11b^+^F480^+^CD206^+^ |

**Table S5. Antibodies list for flow cytometry and immunohistochemistry**

|  |  | **Antibody panel** | **Fluorescence dye** | **Antibody dilution** | **Catalog number** |
| --- | --- | --- | --- | --- | --- |
| Flow cytometry | T-Cells | L/D | Alexa 488 | 400:1 | Invitrogen, #L34970 |
|  |  | CD45^+^ | APC-Cy7 | 400:1 | BD Bioscience, #560501 |
|  |  | TCR^+^ | PE-Cy7 | 200:1 | BD Bioscience, #560729 |
|  |  | CD8^+^ | Pacific blue | 400:1 | BD Bioscience, #558106 |
|  |  | CD4^+^ | BV605 | 400:1 | BD Bioscience, #740336 |
|  |  | Foxp3^+^ | PE | 200:1 | BD Bioscience, #560408 |
|  | MDSC | L/D | Alexa 488 | 400:1 | Invitrogen, #L34970 |
|  |  | CD45^+^ | V450 | 400:1 | BD Bioscience, #560501 |
|  |  | CD11b^+^ | Percp-cy5.5 | 400:1 | BD Bioscience, #550993 |
|  |  | Ly6C^+^ | APC | 200:1 | Biolegend, #128015 |
|  |  | Ly6G^+^ | APC-Cy7 | 200:1 | BD Bioscience, #560600 |
|  | Macrophage | L/D | Alexa 488 | 400:1 | Invitrogen, #L34970 |
|  |  | CD45^+^ | APC-Cy7 | 400:1 | BD Bioscience, #557659 |
|  |  | CD11b^+^ | Percp-cy5.5 | 200:1 | BD Bioscience, #550993 |
|  |  | F4/80^+^ | PE | 200:1 | Biolegend, #123110 |
|  |  | CD86^+^ | BV605 | 200:1 | Biolegend, #105037 |
|  |  | CD206^+^ | PE/Cy7 | 200:1 | Biolegend, #141719 |
| IHC | ICD marker | CRT^+^ | - | 500:1 | Abcam, #ab92516 |
|  |  | HMGB1^+^ | - | 500:1 | Abcam, #ab79823 |
|  | Apoptosis marker | CC-3 | - | 800:1 | Cell Signaling, #9661 |

**Table S6.** **Statistical comparisons of immune-cell populations between groups.** (By one-way ANOVA with Tukey’s test for multiple comparisons (* (p < 0.05), ** (p < 0.01), *** (p < 0.001) (Graphpad Prism, version 7.03). Unpaired t-test, was also performed for the “not significant, ns” groups and is presented in brackets in the cases when the result is different from the one-way ANOVA).

(1) 4T1 model

| Groups | | POx-PLX (75 mg/kg) | POx-PTX (30 mg/kg) | POx-PTX (75 mg/kg) | POx-PTX/PLX  (75 mg/kg PTX and 75 mg/kg PLX3397) |
| --- | --- | --- | --- | --- | --- |
| CD8^+^  T cell | Saline | ns | ** | ** | *** |
|  | POx-PLX (75 mg/kg) |  | *** | *** | *** |
|  | POx-PTX (30 mg/kg) |  |  | ns | ns |
|  | POx-PTX (75 mg/kg) |  |  |  | ns |
| CD4^+^  T cell | Saline | ns | ** | *** | *** |
|  | POx-PLX (75 mg/kg) |  | * | ** | *** |
|  | POx-PTX (30 mg/kg) |  |  | ns | ns (*) |
|  | POx-PTX (75 mg/kg) |  |  |  | ns (*) |
| Treg | Saline | ns (**) | ns (*) | ns (*) | ns |
|  | POx-PLX (75 mg/kg) |  | ** | ** | * |
|  | POx-PTX (30 mg/kg) |  |  | ns | ns |
|  | POx-PTX (75 mg/kg) |  |  |  | ns |
| M-MDSC | Saline | ns | ns (*) | ns | ns |
|  | POx-PLX (75 mg/kg) |  | ns | ns | ns |
|  | POx-PTX (30 mg/kg) |  |  | ns | ns (**) |
|  | POx-PTX (75 mg/kg) |  |  |  | ns (*) |
| G-MDSC | Saline | ns | * | ** | *** |
|  | POx-PLX (75 mg/kg) |  | * | ** | *** |
|  | POx-PTX (30 mg/kg) |  |  | ns | ns |
|  | POx-PTX (75 mg/kg) |  |  |  | ns (*) |
| Macrophage | Saline | *** | ns (*) | * | ns |
|  | POx-PLX (75 mg/kg) |  | *** | *** | *** |
|  | POx-PTX (30 mg/kg) |  |  | ns | ns |
|  | POx-PTX (75 mg/kg) |  |  |  | ns |
| M1  Macrophage | Saline | *** | ns (**) | * | *** |
|  | POx-PLX (75 mg/kg) |  | ns | ns | ns |
|  | POx-PTX (30 mg/kg) |  |  | ns | * |
|  | POx-PTX (75 mg/kg) |  |  |  | ns (***) |
| M2  Macrophage | Saline | ns (*) | ** | * | *** |
|  | POx-PLX (75 mg/kg) |  | ns | ns | *** |
|  | POx-PTX (30 mg/kg) |  |  | ns | ** |
|  | POx-PTX (75 mg/kg) |  |  |  | ** |
| M2/M1  ratio | Saline | ** | * | ** | ** |
|  | POx-PLX (75 mg/kg) |  | ns | ns | ns |
|  | POx-PTX (30 mg/kg) |  |  | ns | ns |
|  | POx-PTX (75 mg/kg) |  |  |  | ns |

(2) T11-apobec model

| Groups | | POx-PLX (75 mg/kg) | POx-PTX (30 mg/kg) | POx-PTX (75 mg/kg) | POx-PTX/PLX  (75 mg/kg PTX and 75 mg/kg PLX3397) |
| --- | --- | --- | --- | --- | --- |
| CD8^+^  T cell | Saline | ns | ns | ns | ns |
|  | POx-PLX (75 mg/kg) |  | ns | ns | ns |
|  | POx-PTX (30 mg/kg) |  |  | ns | ns |
|  | POx-PTX (75 mg/kg) |  |  |  | ns |
| CD4^+^  T cell | Saline | ns | ns | ns | * |
|  | POx-PLX (75 mg/kg) |  | ns (*) | ns (*) | ns (*) |
|  | POx-PTX (30 mg/kg) |  |  | ns | ns |
|  | POx-PTX (75 mg/kg) |  |  |  | ns |
| Treg | Saline | ns | ns | ns | ns |
|  | POx-PLX (75 mg/kg) |  | ns | ns | ns |
|  | POx-PTX (30 mg/kg) |  |  | ns | ns |
|  | POx-PTX (75 mg/kg) |  |  |  | ns |
| M-MDSC | Saline | ns | ns | ns | ns |
|  | POx-PLX (75 mg/kg) |  | ns | ns | ns |
|  | POx-PTX (30 mg/kg) |  |  | ns | ns (*) |
|  | POx-PTX (75 mg/kg) |  |  |  | ns |
| G-MDSC | Saline | ns | ns | ns (*) | ns (**) |
|  | POx-PLX (75 mg/kg) |  | ns | * | ** |
|  | POx-PTX (30 mg/kg) |  |  | ns (*) | ns (*) |
|  | POx-PTX (75 mg/kg) |  |  |  | ns |
| Macrophage | Saline | * | ns | ns | * |
|  | POx-PLX (75 mg/kg) |  | * | ns (*) | ns |
|  | POx-PTX (30 mg/kg) |  |  | ns | * |
|  | POx-PTX (75 mg/kg) |  |  |  | * |
| M1  Macrophage | Saline | *** | ns (*) | ns (*) | ns (**) |
|  | POx-PLX (75 mg/kg) |  | ** | ** | * |
|  | POx-PTX (30 mg/kg) |  |  | ns | ns |
|  | POx-PTX (75 mg/kg) |  |  |  | ns |
| M2  Macrophage | Saline | ns | ns | ns | ns |
|  | POx-PLX (75 mg/kg) |  | ns | ns | ns |
|  | POx-PTX (30 mg/kg) |  |  | ns | ns |
|  | POx-PTX (75 mg/kg) |  |  |  | ns |
| M2/M1  ratio | Saline | *** | ns (*) | ns | * |
|  | POx-PLX (75 mg/kg) |  | * | * | ns (*) |
|  | POx-PTX (30 mg/kg) |  |  | ns | ns |
|  | POx-PTX (75 mg/kg) |  |  |  | ns |

(3) T12 model

| Groups | | POx-PLX (75 mg/kg) | POx-PTX (30 mg/kg) | POx-PTX (75 mg/kg) | POx-PTX/PLX  (75 mg/kg PTX and 75 mg/kg PLX3397) |
| --- | --- | --- | --- | --- | --- |
| CD8^+^  T cell | Saline | ns (**) | ns (*) | ns | ns (**) |
|  | POx-PLX (75 mg/kg) |  | ns | ns | ns (**) |
|  | POx-PTX (30 mg/kg) |  |  | ns | ns |
|  | POx-PTX (75 mg/kg) |  |  |  | ns |
| CD4^+^  T cell | Saline | ns (**) | ns (**) | ** | ns (*) |
|  | POx-PLX (75 mg/kg) |  | ns | * | ns |
|  | POx-PTX (30 mg/kg) |  |  | ns | ns |
|  | POx-PTX (75 mg/kg) |  |  |  | ns |
| Treg | Saline | ns | ns | ns | ns |
|  | POx-PLX (75 mg/kg) |  | ns | ns | ns |
|  | POx-PTX (30 mg/kg) |  |  | ns | ns |
|  | POx-PTX (75 mg/kg) |  |  |  | ns (***) |
| M-MDSC | Saline | *** | ns (*) | ns (***) | ns (***) |
|  | POx-PLX (75 mg/kg) |  | *** | *** | *** |
|  | POx-PTX (30 mg/kg) |  |  | ns | ns |
|  | POx-PTX (75 mg/kg) |  |  |  | ns |
| G-MDSC | Saline | *** | ns | *** | ** |
|  | POx-PLX (75 mg/kg) |  | *** | *** | *** |
|  | POx-PTX (30 mg/kg) |  |  | ns (*) | ns |
|  | POx-PTX (75 mg/kg) |  |  |  | ns |
| Macrophage | Saline | ** | ns (*) | * | * |
|  | POx-PLX (75 mg/kg) |  | *** | *** | *** |
|  | POx-PTX (30 mg/kg) |  |  | ns | ns |
|  | POx-PTX (75 mg/kg) |  |  |  | ns |
| M1  Macrophage | Saline | ns | ns | ns | ns |
|  | POx-PLX (75 mg/kg) |  | ns | ns | ns |
|  | POx-PTX (30 mg/kg) |  |  | ns | ns |
|  | POx-PTX (75 mg/kg) |  |  |  | ns |
| M2  Macrophage | Saline | ns | ns | ns | * |
|  | POx-PLX (75 mg/kg) |  | ns | ns | ns |
|  | POx-PTX (30 mg/kg) |  |  | ns | ns (*) |
|  | POx-PTX (75 mg/kg) |  |  |  | ns (*) |
| M2/M1  ratio | Saline | * | ns | ns | ** |
|  | POx-PLX (75 mg/kg) |  | ns | ns | ns |
|  | POx-PTX (30 mg/kg) |  |  | ns | ns (*) |
|  | POx-PTX (75 mg/kg) |  |  |  | ns (*) |

**Table S7.** **Statistical comparisons of lung metastasis between all groups in Figure 7.** (By one-way ANOVA with Tukey’s test for multiple comparisons (* (p < 0.05), ** (p < 0.01), *** (p < 0.001) (Graphpad Prism, version 7.03). Unpaired t-test, was also performed for the “not significant, ns” groups and is presented in brackets in the cases when the result is different from the one-way ANOVA).

(a) Statistical comparison of lung metastasis in each drug treatment groups

|  | Saline  (Anti-CD4) | Saline  (Anti-CD8) | POx-PTX  (30 mg/kg)  (Anti-CD4) | POx-PTX  (30 mg/kg)  (Anti-CD8) | POx-PTX (75 mg/kg)  (Anti-CD4) | POx-PTX  (75 mg/kg)  (Anti-CD8) | co-loaded POx-PTX/PLX  (75 mg/kg PTX and  75 mg/kg PLX3397)  (Anti-CD4) | co-loaded POx-PTX/PLX  (75 mg/kg PTX and  75 mg/kg PLX3397)  (Anti-CD8) |
| --- | --- | --- | --- | --- | --- | --- | --- | --- |
| Saline (No antibody) | ns | ** | ns | ns (*) | ns | ns | ns | ns |
| Saline (Anti-CD4) | - | ns (**) | ns | ns (**) | ns | ns | ns | ns |
| POx-PTX (30 mg/kg)  (No antibody) | ns | ns | ns | ns | ns | ns | ns | ns |
| POx-PTX (30 mg/kg)  (Anti-CD4) | ns | ns (*) | - | ns | ns | ns | ns | ns |
| POx-PTX (75 mg/kg)  (No antibody) | ns (*) | *** | ns | ** | ns | ns | ns (*) | ns (*) |
| POx-PTX (75 mg/kg)  (Anti-CD4) | ns | * | ns | ns | - | ns | ns | ns |
| co-loaded POx-PTX/PLX  (75 mg/kg PTX and 75 mg/kg PLX3397) (No antibody) | ns (**) | *** | ns (*) | *** | ns | ns (*) | ns (**) | ns (***) |
| co-loaded POx-PTX/PLX  (75 mg/kg PTX and 75 mg/kg PLX3397) (Anti-CD4) | ns | * | ns | ns (***) | ns | ns | - | ns |

(b) Statistical comparison of lung metastasis in each antibody treatment groups

|  | | POx-PTX (30 mg/kg) | POx-PTX (75 mg/kg) | co-loaded POx-PTX/PLX  (75 mg/kg PTX and 75 mg/kg PLX3397) |
| --- | --- | --- | --- | --- |
| No antibody | Saline | ns | ns | * |
|  | POx-PTX (30 mg/kg) |  | * | ** |
|  | POx-PTX (75 mg/kg) |  |  | ns |
| Anti-CD4  (CD4^+^ depletion) | Saline | ns | ns | ns |
|  | POx-PTX (30 mg/kg) |  | ns | ns |
|  | POx-PTX (75 mg/kg) |  |  | ns |
| Anti-CD8  (CD8^+^ depletion) | Saline | ns | * | ** |
|  | POx-PTX (30 mg/kg) |  | * | * |
|  | POx-PTX (75 mg/kg) |  |  | ns |
